# Pumping the brakes on RNA velocity – understanding and interpreting RNA velocity estimates

**DOI:** 10.1101/2022.06.19.494717

**Authors:** Shijie C. Zheng, Genevieve Stein-O’Brien, Leandros Boukas, Loyal A. Goff, Kasper D. Hansen

## Abstract

RNA velocity analysis of single cells promises to predict temporal dynamics from gene expression. Indeed, in many systems, it has been observed that RNA velocity produces a vector field that qualitatively reflects known features of the system. Despite this observation, the limitations of RNA velocity estimates are poorly understood. Using real data and simulations, we dissect the impact of different steps in the RNA velocity workflow on the estimated vector field. We find that the process of mapping RNA velocity estimates into a low-dimensional representation, such as those produced by UMAP, has a large impact on the result. The RNA velocity vector field strongly depends on the k-NN graph of the data. This dependence leads to significant estimator errors when the k-NN graph is not a faithful representation of the true data structure, a feature that cannot be known for most real datasets. Finally, we establish that RNA velocity estimates expression speed neither at the gene nor cellular level. We propose that RNA velocity is best considered a smoothed interpolation of the observed k-NN structure, as opposed to an extrapolation of future cellular states, and that the use of RNA velocity as a validation of latent space embedding structures is circular.

## INTRODUCTION

RNA velocity analysis is widely used to infer temporal dynamics in single-cell gene expression data. In it’s original definition, RNA velocity is the time derivative of gene expression state (*ds/dt* with *s* representing the high-dimensional expression state and *t* time)(La Manno et al., 2018). Based on earlier work of Zeisel et al. (2011), La Manno et al. (2018) propose a model built using an ordinary differential equation model of transcription and the assumption that the relationship between spliced mRNA expression and unspliced pre-mRNA expression can be used to infer whether a gene is in the process of being up- or down-regulated or in a steady expression state. Using this model, they derive an estimator for velocity using single-cell expression data. This model and estimation procedure was later extended by Bergen et al. (2020). These two variants are known as the steady-state model (La Manno et al., 2018) and the dynamical model (Bergen et al., 2020).

The primary output of an RNA velocity analysis for single-cell gene expression data is a vector field visualized on a low-dimensional embedding, usually constructed using UMAP or t-SNE. This vector field is supposed to represent local changes in expression state; one might describe the output as an alternative to trajectory inference analysis (Weiler et al., 2021). This output is often used to provide evidence for the correctness of the embedding. An intermediate output towards the low-dimensional vector field is the high-dimensional estimation of *ds*/*dt*; we refer to these estimates as “velocities” (or gene-specific rates of change). This estimation is performed gene-by-gene using the biophysical ordinary differential equations. We will use “velocity vector field” to refer to the low-dimensional vector field obtained by mapping the velocities into a low-dimensional space. In many datasets, the visualization of the velocity vector field appears to reflect what is known about the biological system and the relationship between cell states/types in the system.

Despite the widespread use and popularity of RNA velocity, studies that validate the highdimensional RNA velocity estimates at the gene level are lacking. A significant reason is that measuring the instantaneous rate of expression change in a single cell is extremely hard. Q Qiu et al. (2020) used metabolic labeling coupled with scRNA-seq to distinguish between newly synthesized mRNA and older mRNA. They directly compared the velocity estimates from metabolic labeling with the splicing-kinetics based RNA velocity of 3 genes and found a poor correspondence between the two types of velocity estimates. It can be argued, however, that the explicit aspects of mRNA biogenesis estimated by metabolic labeling are subtly different from the models used in RNA velocity.

The potential of predicting the future state of individual cells has spurred tremendous interest in RNA velocity among the single-cell research community, resulting in many reviews and work building off the velocity framework. On the review side, Bergen et al. (2021) highlights examples where the RNA velocity vector is not compatible with the known biology of the system and proposes that this is due to assumptions of multiple kinetic regimes, transcriptional boosts, high noise, or time-constant rates of transcription, splicing, and degradation. They further envision using gene regulatory networks and multimodal omics to expand RNA velocity models. Gorin et al. (2022) is an in-depth discussion and critique of RNA velocity and complements the work we present here. The paper covers the underlying mathematics of RNA velocity in detail and advocates for a more rigorous approach to RNA velocity, respecting the discrete nature of transcription in single cells. Although the work covers all aspects of RNA velocity, there is a particular focus on the inherent discrete nature of the problem and its implications for preprocessing and biophysical modeling. The manuscript primarily discusses the python implementation of velocyto.

While numerous tools seek to improve the RNA velocity framework, many methods build directly off the original framework. For example, Dynamo integrates metabolic labeling into the splicing-unspliced-dynamics-based model and tries to mathematically recover the whole velocity vector field in the low-dimensional embedding even for regions without any cells existing (X Qiu et al., 2022). Cell-Rank combines the k-NN graph built from the expression profile with the RNA velocity transition probability to detect initial, intermediate, and terminal populations during differentiation (Lange et al., 2022). Marot-Lassauzaie et al. (2022) first discusses several theoretical and computational problems and then proposes two alternative RNA velocity or RNA velocity flavored approaches. First, *κ*-velo tries to address the scale invariance problem by incorporating cell densities with the assumption that cell density is inversely proportional to the average velocity for each gene. Second, eco-velo is a heuristic RNA velocity flavored approach that does not infer high-dimensional RNA velocity. Instead, eco-velo uses the first mutual nearest neighbor (MNN) of a cell in the unspliced matrix space to the spliced matrix space as the proxy for the future state in the spliced matrix space. A vector field is created based on the displacements between the future state and the current state in the spliced matrix space. Gao et al. (2022) proposes a revised high-dimensional RNA velocities estimation model UniTVelo which imposes a gene-shared cell latent time to circumvent the independent estimation issue of other RNA velocity estimations approaches.

Here, we deconstruct the underlying workflow by separating the (gene-level) velocity estimation from the vector field visualization. We then analyze how the methods for mapping and visualizing the vector field impact the interpretation of RNA velocity and discover the central role played by the k-NN graph in both velocity estimation and vector field visualization. Using both simulations and real data, we identify situations where RNA velocity estimates are accurate and evaluate the extent to which the visualizations allow us to discover new structures in the data. We also explore whether – as their name suggests – velocity estimates can provide quantitative information about the speed at which cells progress along a trajectory.

## RESULTS

### RNA velocity analysis and its implementations

RNA velocity analysis consists of two primary steps:

1. Spliced and unspliced counts are preprocessed (smoothed/imputed), and a cell-specific velocity is estimated separately for each gene.
2. The high-dimensional velocity estimates are mapped into a low-dimensional embedding, and the resulting vector field is visualized on this embedding.

Assessments of RNA velocity analysis results are usually qualitative: does the resulting vector field visualization accurately capture known cellular dynamics of the biological system? This endeavor centers on the resulting vector field visualization (step 2), and we will discuss its properties in detail.

The two primary approaches to RNA velocity are the steady-state model (La Manno et al., 2018) and the dynamical model (Bergen et al., 2020). The main differences between the two are the specific assumptions about the parameters in the biophysical models of transcript abundance that are used to estimate cell-specific velocities (step 1 above), although many other differences impact the result (Supplementary Note 3.1). Here, we primarily focus on the newer dynamic model (Bergen et al., 2020), as implemented in scVelo. We also occasionally make comparisons to the steady-state model, also using its scVelo implementation (which we chose over other implementations of the steady-state model to ensure that we can keep other parameters of the workflow constant)

The complete series of operations involved in an RNA velocity analysis workflow is outlined in Figure 1 using the specific implementation for scVelo (note that Gorin et al. (2022) has a similar velocyto-centric figure). We divide Step 1 (inference of high-dimensional RNA velocity values) into two parts: preprocessing and gene-level velocity estimation (Figure 1a). In the preprocessing step, the raw count data is smoothed by a k-NN graph constructed exclusively from the spliced counts. A number of additional steps, such as library size adjustment and log-transformation, are performed and discussed in Gorin et al. (2022). Following this, cell-specific velocities are estimated by fitting the smoothed spliced and unspliced counts to the appropriate biophysical model described by either the steady-state or dynamical model for each gene independently (Methods). Importantly, valid velocity estimations are only retained from a reduced set of genes where the model is considered well fit.

**Figure 1.**
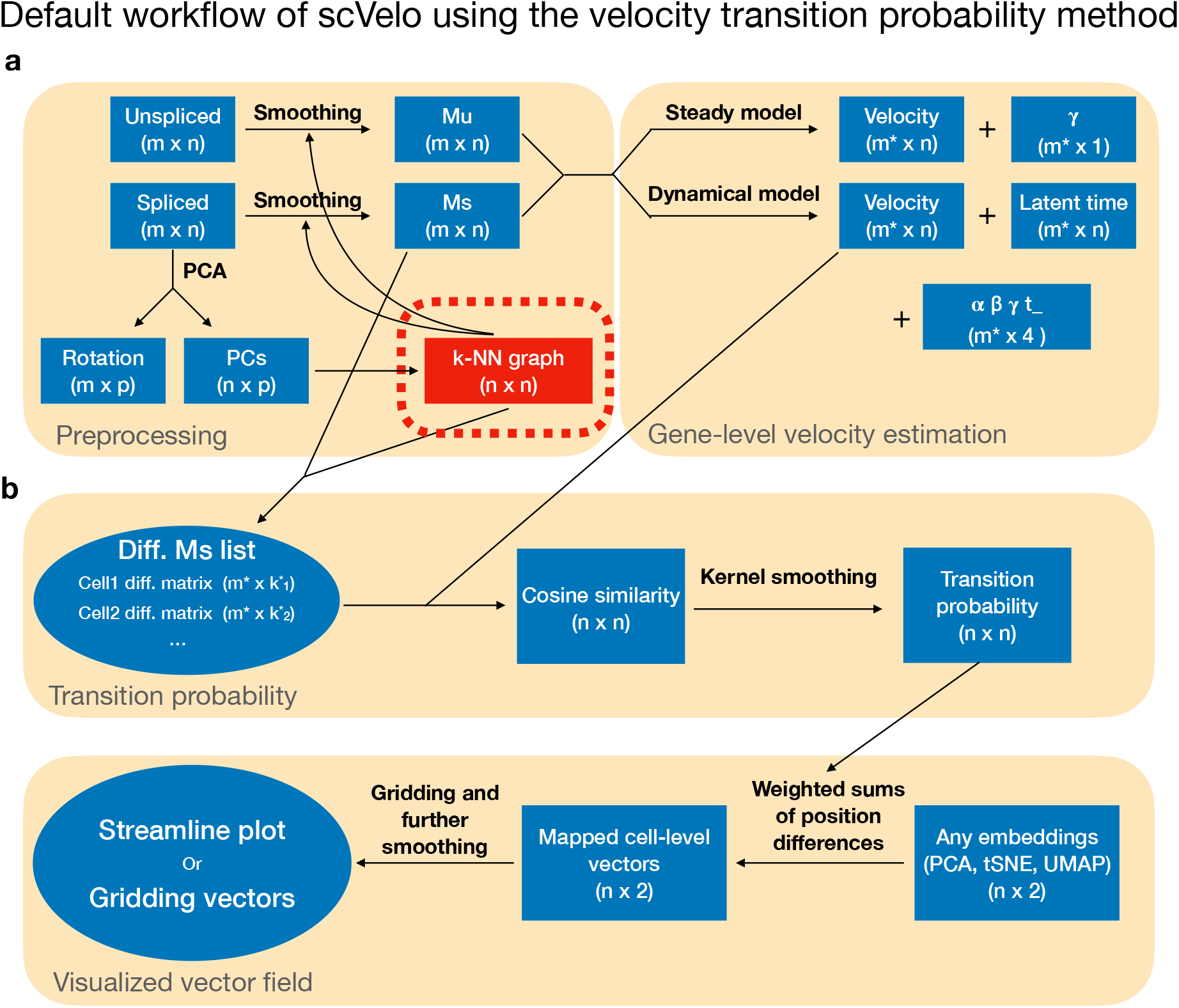
The flowchart of RNA velocity implementation in scRNA-seq data. The graph reflects scVelo. **(a)** A k-NN graph constructed from a PCA of the spliced counts is used to smooth (impute) the spliced and unspliced count matrices, resulting in the Ms and Mu matrix. This is followed by gene-specific velocity estimation using either the dynamical or the steady-state model. **(b)** Visualization of the estimated velocities on a low-dimensional embedding using velocity transition probabilities. First, transition probabilities are computed by considering which neighbors have a difference between the expression of the neighbor and the expression of the cell in question most similar to the estimated velocities. These transition probabilities are used to compute a vector as a linear combination of existing displacements. Finally, the resulting vector field can be visualized using streamline plots or a gridding approach.

In the second step, after estimating the highdimensional velocities, they are then visualized^1^ on a low-dimensional embedding, usually constructed using UMAP or t-SNE from principal components learned from the spliced count matrix. This consists of two steps (Figure 1b). First, the velocity estimates are mapped into the existing lowdimensional embedding. The most common solution is to represent the velocities in this reduced dimensional space using transition probabilities, a method developed by La Manno et al. (2018) and further modified by Bergen et al. (2020). After mapping, we are left with low-dimensional velocity vectors that are visualized using approaches such as streamline plots or gridding average vectors (Supplementary Note3.2). In practice, this last visualization step almost always includes an additional smoothing, such as kernel smoothing over the embedding; furthermore, the lengths of summarized vectors are usually rescaled.

Most attention in the literature has been given to a qualitative visual assessment of the RNA velocity vector field. We, therefore, start by examining how parts of the workflow impact visualization.

### Mapping velocities into a low-dimensional embedding

To visualize scRNA-seq data on a nonparametric embedding, such as t-SNE or UMAP, mapping velocity estimates into an existing embedding is a nontrivial problem (unlike recomputing a new embedding based on the expression data supplemented with the predictions), and we briefly describe existing approaches to this problem. La Manno et al. (2018) introduced a method we will refer to as “velocity transition probabilities” (later modified by Bergen et al. (2020) which is the version we focus on here). This method is central to RNA velocity analysis since it is used in most RNA velocity vector field visualizations. The method also serves as the basis for the CellRank method for predicting fate specification (Lange et al., 2022). Velocity transition probabilities are claimed to provide a solution to the mapping problem, which is compatible with *any* type of low dimensional embedding, including UMAP, t-SNE, and PCA, provided the embedding is constructed using expression data from the same cells yielding the velocity estimates. In addition to velocity transition probabilities, we have embedding-specific approaches such as an orthogonal projection operator for principal component plots and UMAP-transform for mapping into UMAP space (a method supplied by the UMAP authors) (McInnes, 2021).

After estimating high-dimensional velocity by either the steady-state or dynamical model, we have the observed current expression state *s* and the estimated velocity *v* (both high-dimensional) (Methods). A first-order Taylor expansion of the expression state yields the following approximate relationship between current expression level *s*, velocity *v* = *ds/dt* and future expression level *s*^*^:

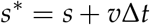

which requires a choice of time step Δ*t*. For a fixed Δ*t*, there is a one-to-one relationship between the velocity and the future expression.

If we can compute *E*(*y*) where *y* is any point in the input state, we can map velocities as

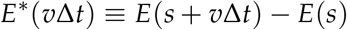

Here we are using *E*^*^ to indicate that the left-hand side is a new operator defined by the right-hand side. If the mapping operator is linear (which is the case for principal component analysis), we get

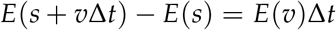

and there is no difference between *E*\*(*v*) and *E*(*v*), and the impact of choosing a time step Δ*t* is just an overall scaling. If the mapping operator is not linear (which we believe is the case for UMAP-transform), there is a difference between *E*^*^(*v*) and *E*(*v*), and the time step matters.

These computations require us to be able to compute *E*(*y*) where *y* is an arbitrary new point in the expression space. The velocity transition probabilities were designed to work in cases where this is not directly obvious, as in the case of nonlinear embeddings. The only points we have available in the embedding space are the mapping of the observed cells, denoted as *E*(*s*_1_),…, *E*(*s_n_*). If we focus on a specific cell *i*, the idea is to represent the velocyto embedding as a weighted sum of empirical differences:

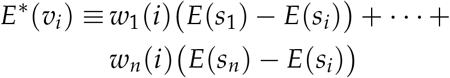

for suitable choices of weights (depending on the cell in question). The weights are constructed to give higher weights to cells with a higher cosine similarity between *s_j_* – *s_i_* and *v_i_* (an approach that bypasses the choice of a time step Δ*t*). In other words, you compute a future expression state in which high weights are given to cells with closer neighbors on the k-NN and to which you believe the cell is transitioning. In scVelo, this sum is restricted to include only terms from neighbors of neighbors of the cell *i* in the k-NN graph of the spliced counts. A consequence of this approach is that the direction of the embedded velocity must be towards the (convex hull) of neighbors of neighbors of the cell. This has the undesirable effect of assuring that the direction of the velocity vector for a given cell is entirely dependent on the expression states of its nearest neighbors, with consequences of constraining the vectors pointing to other existing cells around the query cell (detailed below).

### Transition probabilities impose directional constraints

To illustrate the critical influence of neighboring cells on the resulting vector field, we consider the pancreas dataset featured in Bergen et al. (2020) using the scVelo package for velocity estimates (Figure 2a, b). As previously described, a standard velocity analysis suggests that pre-endocrine cells (orange) flow towards beta cells (light green) as expected. To demonstrate the dependence on nearest neighbor expression estimates, we then fix the highdimensional velocity estimates (which are based on the full dataset) but remove the beta cells (light green) from the embedding step. Because we remove the beta cells, the recomputed velocity transition probabilities force the vectors to point to another part of the available embedding: the alpha cells (dark blue) (Figure 2c and Supplementary Figure S1a), resulting in a dramatic difference in the interpretation of the future state of the pre-endocrine cells. Figure 2d is a quantitative display of the change in low-dimensional vector directions for all cell-specific vectors of pre-endocrine cells in the red rectangle as a result of removing the beta cells from the embedding.

**Figure 2.**
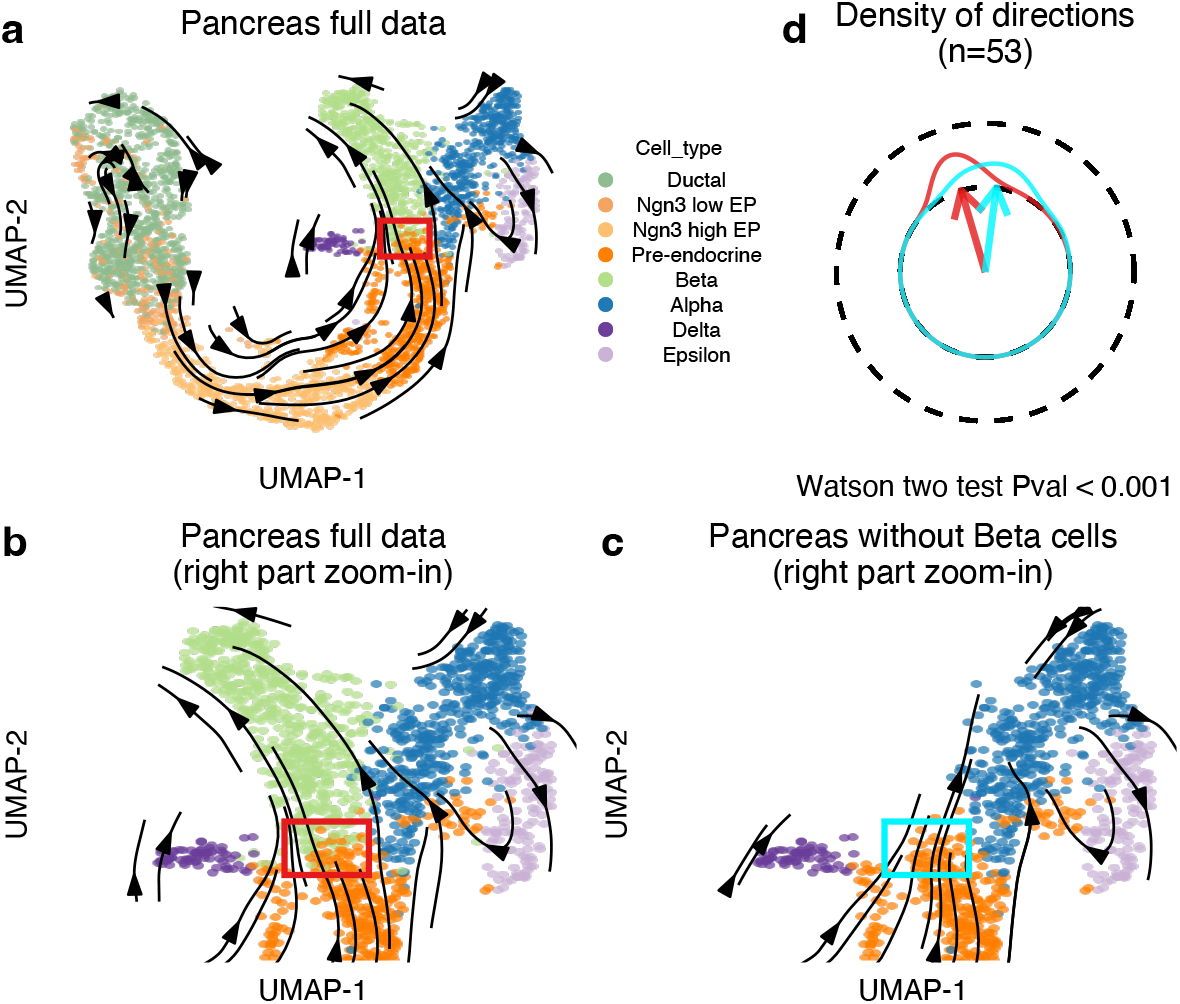
Embedded RNA velocity of the pancreas dataset. **(a)** Dynamical model based RNA velocity embedded using velocity transition probability method for the full pancreas dataset. **(b)** Zoom-in of the right part of (a). **(c)** As (b), but now we show the zoom-in for the pancreas data after removing the beta cells. Note that the gene-level velocity estimations are not changed, but the transition probability matrix is re-computed. **(d)** Comparison of directions of the cell-level vectors for the same set of pre-endocrine cells in the rectangle in (b) and (c).

This example illustrates a significant caveat of using transition probabilities for the low-dimensional estimation of velocity vectors. Velocity transition probabilities cannot represent unseen parts of an expression state space. Importantly, this suggests that the visualization of RNA velocities is more akin to an interpolation of observed expression states and less of an extrapolation of the future states. An example of this impacting data analysis is the comparison of multiple samples with sample-specific cell types or states, perhaps as a consequence of spatial heterogeneity (i.e., technical). In this case, the missing cell types or states will locally warp the vector field using transition probabilities.

### Visualizing velocity without noise

To further examine the impact of velocity transition probabilities, we next employ a simulation experiment in which we control and account for varying noise levels. We follow the simulation strategy of Bergen et al. (2021) and generate 500 cells with 10 genes following the dynamic model with a low noise level. Figure 3a depicts one of these genes and shows that the dynamics of the simulated gene fit well with the dynamical model. To focus on visualization instead of high-dimensional velocity estimation, we use our simulation to define true velocities (defined as the velocity of the expression state prior to adding random noise). Our simulated data have a single trajectory without cycles or bifurcations, which is captured by our UMAP representation (Figure 3b, c). Using velocity transition probabilities for visualization, we obtain a vector field that flows in the correct direction (color indicates time). There is one issue with this vector field: At the start of the field, all vectors are close to zero with a seemingly random direction; this is not supported by the simulations, as the true velocities of these cells are relatively uniform in their direction and magnitude. In contrast, using UMAP-transform for visualization, we obtain a vector field largely similar to the one created using velocity transition probabilities, except around the start of the trajectory, where we observe substantially longer vectors with the correct direction.

**Figure 3.**
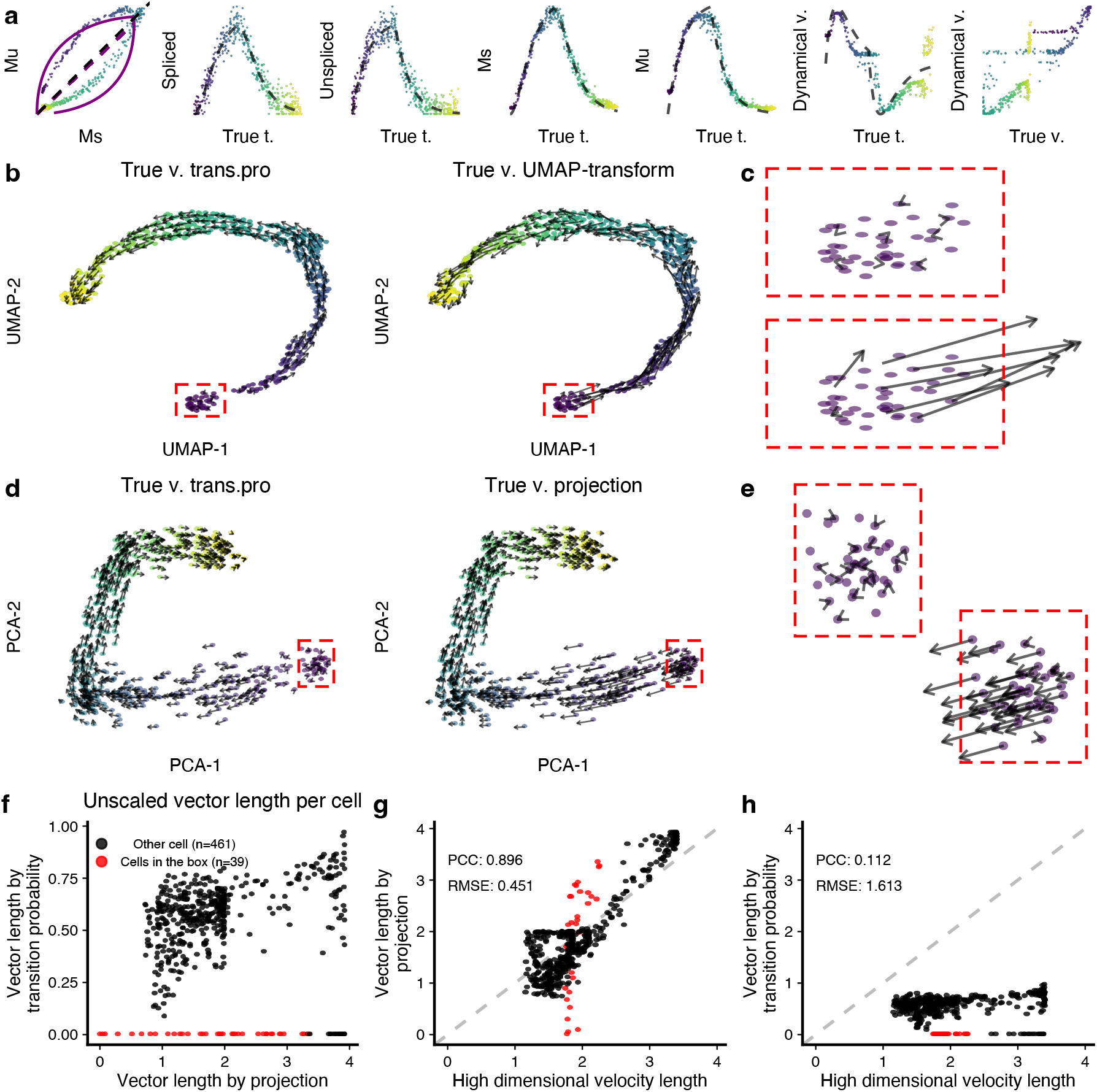
The challenges in visualizing RNA velocity in embeddings. **(a)** Example of a gene in the simulated data. From the left to the right, we show the phase portrait, spliced counts over true latent time, unspliced counts over true latent time, Ms over true latent time, Mu over true latent time, estimated velocity over true latent time, comparisons of estimated velocity and true velocity. **(b)** True RNA velocity vector field on UMAP using the transition probability method (left sub-panel) and UMAP-transform method (right sub-panel). **(c)** Zoom-ins of two red rectangle labelled regions in (b). **(d)** As (b), but on PCA. **(e)** Zoom-ins of two red rectangle labelled regions in (d). **(f)** Comparison of the cell-level vector length produced by two methods in (e). **(g)** Comparison of the cell-level vector length between the high-dimensional vector (gene-level velocity) and the low-dimensional vector produced by PCA projection. **(h)** Comparison of the cell-level vector length between the high-dimensional vector (gene-level velocity) and the low-dimensional vector mapped by the transition probability method.

We next assessed the impact of velocity transition probabilities when mapping velocities to a linear embedding (PCA). For our simulated data, principal component analysis reveals the expected single trajectory in the first two dimensions with a topology that is highly similar to that of the UMAP plot (Figure 3d, e). The median of cosine similarities of cell-level vectors produced by transition probability and direct projection is as high as 0.984, representing a high level of agreement in directions. However, in this linear space as well, the velocity transition probabilities have issues correctly mapping the vectors around the start of the trajectory. Using the PCA projection operator, which can be considered the truth in this particular embedding, reveals vectors of substantial length and the correct direction.

To bypass any kind of visualization summarization that might mask local discrepancies, we compare the vector lengths obtained by the two approaches for each cell and confirm that the start of the trajectory is an area of high discordance between the two mapping methods (Figure 3f). Furthermore, we can quantify the relationship between high-dimensional velocities (true velocities in this simulation) and embedded velocities. This reveals the length of the PCA-projected velocities to be related to the length of the true high-dimensional velocities (Figure 3g). In contrast, the lengths of the embedded vectors using velocity transition probabilities do not have any meaningful relationship with the high-dimensional vectors (Figure 3h). Note that, in physics, speed is defined as the length of the velocity vector.

In conclusion, in this simulation experiment, the different mapping approaches yield a qualitatively similar vector field with an exception around the start of the trajectory where the velocity transition probabilities fail. Notably, the reasons for why this particular portion of the transition probability mapping fails are unclear, but this failure is not observed when using ‘projection-based’ methods for low-dimensional velocity vector field mapping. Furthermore, for all simulated cells, there is little to no relationship between the length of the vectors mapped using velocity transition probabilities and the true high-dimensional velocities. Using a PCA-based projection does a substantially better job at reflecting velocity vector lengths in high-dimensional space.

### Velocity estimation and visualization are strongly dependent on the k-NN graph

Single-cell data are known to be noisy. We next asked how increasing noise might affect lowdimensional vector field visualization. In the previous section, we examined issues with vector field visualization in a simulation setting with (unrealistic) low noise. When we use the same dynamics but increase the noise by 5x, the resulting PCA changes from a one-dimensional manifold to a ball (Figure 4a). Different time points are located roughly in distinct quadrants of the ball and the time progression is clockwise. A similar ball-like observation is made when we visualize the data using UMAP (Supplementary Figure S2). Again, we project the true velocities using the PCA projection matrix, and the resulting vector field broadly reflects the time progression (as expected) (Figure 4b). Note that neighboring vectors belonging to different quadrants point in very different directions, giving the impression of a very noisy visualization unless the true time progression is already known for reference. Notably, using a visualization method such as a streamline plot will have a smoothing effect that hides this behavior. We consider this vector field to be the gold standard for this embedding.

**Figure 4.**
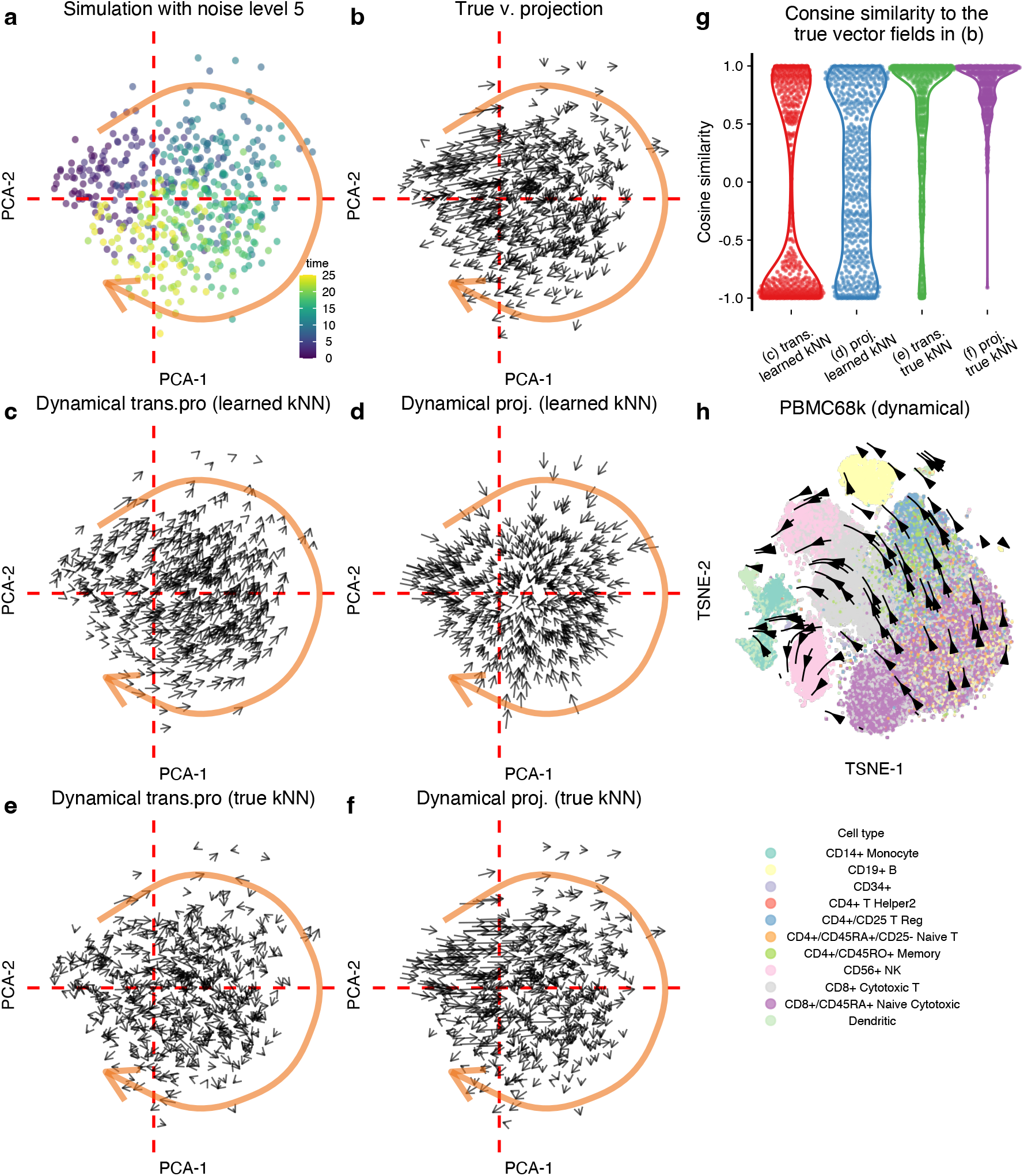
The RNA velocity vector field is dependent on the k-NN graph. **(a)** PCA of the simulated data. The dashed red line is an auxiliary line for mapping locations in other panels. **(b)** The PCA projections of the true RNA velocities. **(c)** RNA velocity inferred using the dynamic model and visualized using velocity transition probabilities. **(d)** RNA velocity inferred using the dynamic model and visualized using PCA projection. **(e)** As (c), we use the true k-NN graph to both preprocess the data and as input to the velocity transition probabilities. **(f)** As (d), but we use the true k-NN graph to preprocess the data. **(g)** The cosine similarities between the mapped cell-level vectors in (c)-(f) and the “true” mapped cell-level vectors in (b). The usage of true k-NN substantially improves the cell-level vectors for both transition probability and direct projection methods. **(h)** The RNA velocity vector field of PBMC68k data flows with the embeddings, but it does not reflect true biological trajectory. Each point represents a cell, which is colored by cell type.

If we estimate velocities using the dynamical model and use the velocity transition probability method for visualization, we obtain a vector field with a smooth progression from left to upper right (Figure 4c). Although visually pleasing (with apparent consistency in direction amongst neighbors), more than 50% of the vectors are strongly dissimilar from the gold standard (Figure 4g). Note that this comparison is made at the cell level; the visualizations in Figure 4 use the gridding approach. If we keep the velocity estimates from the dynamical model but map them using the PCA projection, we obtain a flow going from the edge of the circle towards its center (Figure 4d); not reflecting the true time progression. Because neither visualization method compares favorably to the gold standard, it is tempting to conclude that this is a result of noise overwhelming the velocity estimates (we observe similar failures for the steady-state model in the Supplementary Figure S3).

However, using the right k-NN graph is critical for correct inference. When we increase the noise level, two things are happening simultaneously. Gene-level measurements have added noise, and the k-NN graph learned from the data is perturbed away from the true graph; the latter is reflected by the significant change in the PCA layout. To investigate why the vector fields are wrong, we obtain the true k-NN graph and use this k-NN graph to preprocess both the spliced and unspliced matrices and to construct the velocity transition probabilities. Using the true k-NN graph yields a vector field that is substantially more aligned to the gold standard; there is a slight improvement when using the PCA projection matrix compared to velocity transition probabilities (Figures 4e-g). The gridding visualization of the velocity transition probabilities gives a noisy impression, which is not fully reflected by the cell-level comparisons in Figure 4g.

Is the improvement in using the true k-NN graph driven by its usage in preprocessing or in mapping? When we attempt combinations of true and learned k-NN graphs for the two steps, we observe the most notable improvement by using the true k-NN graph in the preprocessing step (Supplementary Figure S3).

We draw several conclusions from this example. First, it reveals that RNA velocity is critically dependent on the k-NN graph for vector field visualization. Second, we see how the true k-NN graph can be distorted by noise, an observation that is likely to be relevant outside of RNA velocity, given the importance of k-NN graphs in single-cell expression analysis. Third, we observe that the smoothness of the vector field does not imply correctness. Fourth, we are intrigued by the similarity of the “ball-of-cells”-like embedding to some existing single-cell embeddings: we refer to embeddings that show a dense structure where different parts of the structure appear to consist of distinct cell types. An example is the data depicted in Figure 4h, which is comprehensively discussed in Bergen et al. (2021) as an example where the RNA velocity fails. Our simulation experiment suggests this could arise primarily from noise deforming the k-NN graph.

### Evaluating gene-level RNA velocities using simulations

So far, we have focused on the low-dimensional velocity vector field on a fixed embedding. It is natural to ask to what extent the gene-level estimates are accurate. To do so, we turn to our simulations. Using a noise level of 3, the topology of the PCA plot is a 1-dimensional trajectory (Supplementary Figure S4) with some noise. Using a k-NN learned from the data, the estimated velocities are quite inaccurate when considered as a function of the true time (Figure 5a-c, Supplementary Figure S5 for additional quantities). When switching to the true k-NN graph, the dynamical model works substantially better, although there is still some discrepancy between the true velocity and the estimated velocity (Figure 5d-f). Supplementary Figure S5 expands on the quantities depicted in Figure 5 and Supplementary Figure S6 depicts the situation when we increase the noise level to 5, the situation where the PCA plot changes from a 1-dimensional trajectory to a ball.

**Figure 5.**
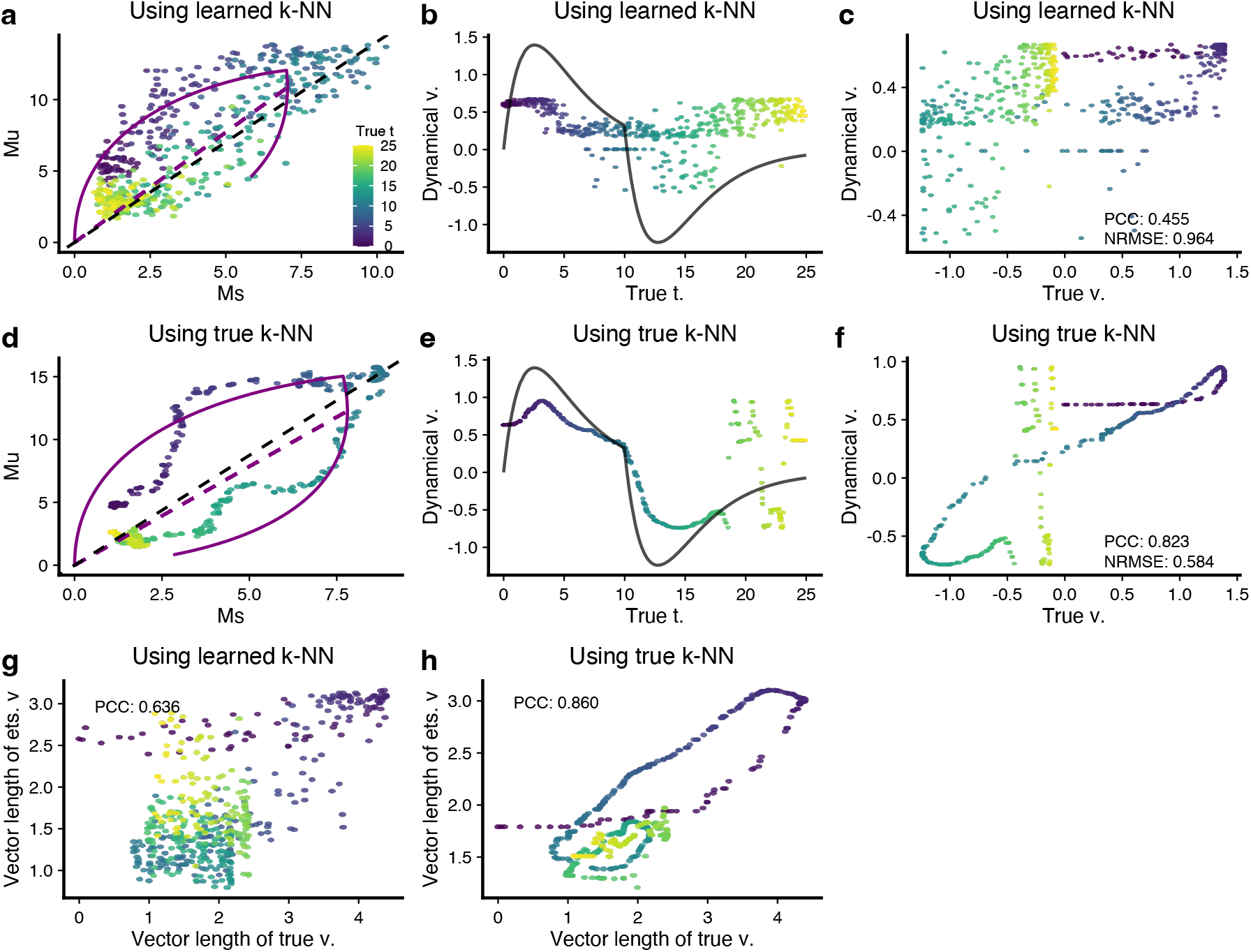
Gene-level RNA velocity estimation depends on the underlying k-NN graph. In all panels, a data point represents a cell and is colored by the known true latent time *t*. All black solid lines represent the know true values. **(a)** Phase portrait shows the Ms over Mu using the learned k-NN. The dynamics are estimated by the dynamical model. **(b)** Estimated velocity (points) using the learned k-NN and true velocity (black line) over true latent time *t*. **(c)** Scatter plot compares the estimated velocity values (using the learned k-NN) to the true velocity values. Pearson correlation coefficient (PCC) and Normalized Root Mean Square Error (NRMSE) are given. **(d-f)** As (a-c), but now we use the true k-NN to get Ms and Mu matrices. The estimated velocity values are much closer to the true velocity values with PCC 0.823 and NRMSE 0.584. **(g)** Scatter plot compares the vector length of true (high-dimensional) velocity to that of the estimated (high-dimensional) velocity by the dynamical model using the learned k-NN graph. **(h)** As (g), but we use the true k-NN to infer (high-dimensional) velocity.

Supplementary Figure S7a, b depicts a comprehensive evaluation across all genes and across many noise settings for both the dynamical and the steady-state model. As the noise level increases, the dynamical model cannot be fitted to most of the genes, and the performance measures are restricted to the few genes that provide a passable fit (as determined by the software). We make several observations. First, the performance is markedly better when using the true k-NN graph compared to the learned k-NN graph. Second, the steady-state model performs better when the assessment criteria are independent of scale (such as PCC), whereas the dynamical model performs better when the assessment criteria are scale-dependent (such as NRMSE).

Aside from comparing the velocity values gene-by-gene, we can compare the speed and direction of the high-dimensional velocity vector cell-by-cell. Speed is the length of the velocity vector. Using a noise level of 3, we find poor concordance between the estimated true speed when using the learned k-NN (Figure 5g). This is substantially improved by using the true k-NN (Figure 5h). The steadystate model shows a similar behavior (Supplementary Figure S8), although the scale of the length is wrong. We can compare the speed estimates with the truth using both absolute differences (a measure dominated by the scale of the length) and Pearson correlation (unaffected by the scale) (Supplementary Figure S9a,b). Together, these two measures reveal that the estimates of speed from both the dynamical and the steady-state model fail to reflect the truth, at least when the learned k-NN graph is used. Direction is harder to assess in high dimensions; the cosine similarity is a step toward this goal, and it suggests poor concordance between estimated and true directions when using the learned k-NN graph (Supplementary Figure S9c). Using the true k-NN graph leads to substantial performance improvements in estimating both speed and direction, but the overall performance is still poor for high noise levels.

In conclusion, the gene level RNA velocity estimations are highly dependent on the k-NN graph used to smooth the data. Even in a simulated setting where the observed k-NN graph reflects the true underlying structure (the PCA plot shows a 1-dimensional trajectory), using the observed k-NN graph results in substantial errors in estimated velocities.

### Correct estimate of the vector field does not imply accurate high-dimensional velocity estimation

Our ability to assess RNA velocity is limited by the technical difficulties in measuring the instantaneous rate of change in expression for many genes in single cells. To evaluate the RNA velocity estimates in real data, we consider a recent dataset on the cell cycle measured using the FUCCI system combined with scRNA-seq (Mahdessian et al., 2021). The cell cycle is a well-understood periodic process (Whitfield et al., 2002), where cell cycle-related genes go through phases of up- and downregulation. In this dataset, we can place each cell in a continuum representing the cell cycle. One approach to this goal is to take advantage of the FUCCI system, which tracks cell cycle time by the protein levels of two key cell cycle regulators, CDT1 and GEMININ. However, in our recent work, we have established that it is possible to improve this cell cycle time by projecting this data into a lowdimensional space representing the cell cycle (SC Zheng et al., 2022). We refer to these two time representations as the tricycle cell cycle time (position) and the FUCCI pseudotime.

First, we note that cell cycle genes have a specific role in RNA velocity estimation. Due to the periodicity, cell cycle genes go through both an up- and a down-regulation phase, but, perhaps surprisingly, they dominate among such genes in existing singlecell datasets. In practice, this suggests that dynamically regulated cell cycle genes *should* sample both the increasing and decreasing portions of the RNA velocity biophysical models, providing for a potential better fitting of these models. Indeed, if we look at the top ten best fitting genes across ten different single-cell expression datasets, the *only* genes having a “complete” phase portrait are cell cycle genes (Supplementary Figure S10). Non-cell-cycle genes are either in the induction or repression phase. For this reason, we believe that cell cycle datasets such as the FUCCI dataset we consider here are amongst the most suitable for RNA velocity analysis using velocity estimates derived from current biophysical models.

Using the FUCCI data, the visualized vector field reflects cell cycle time (Figure 6a), and so does the phase plot of the 10 best fitting cell cycle genes (Figure 6b and Supplementary Figure S11).

**Figure 6.**
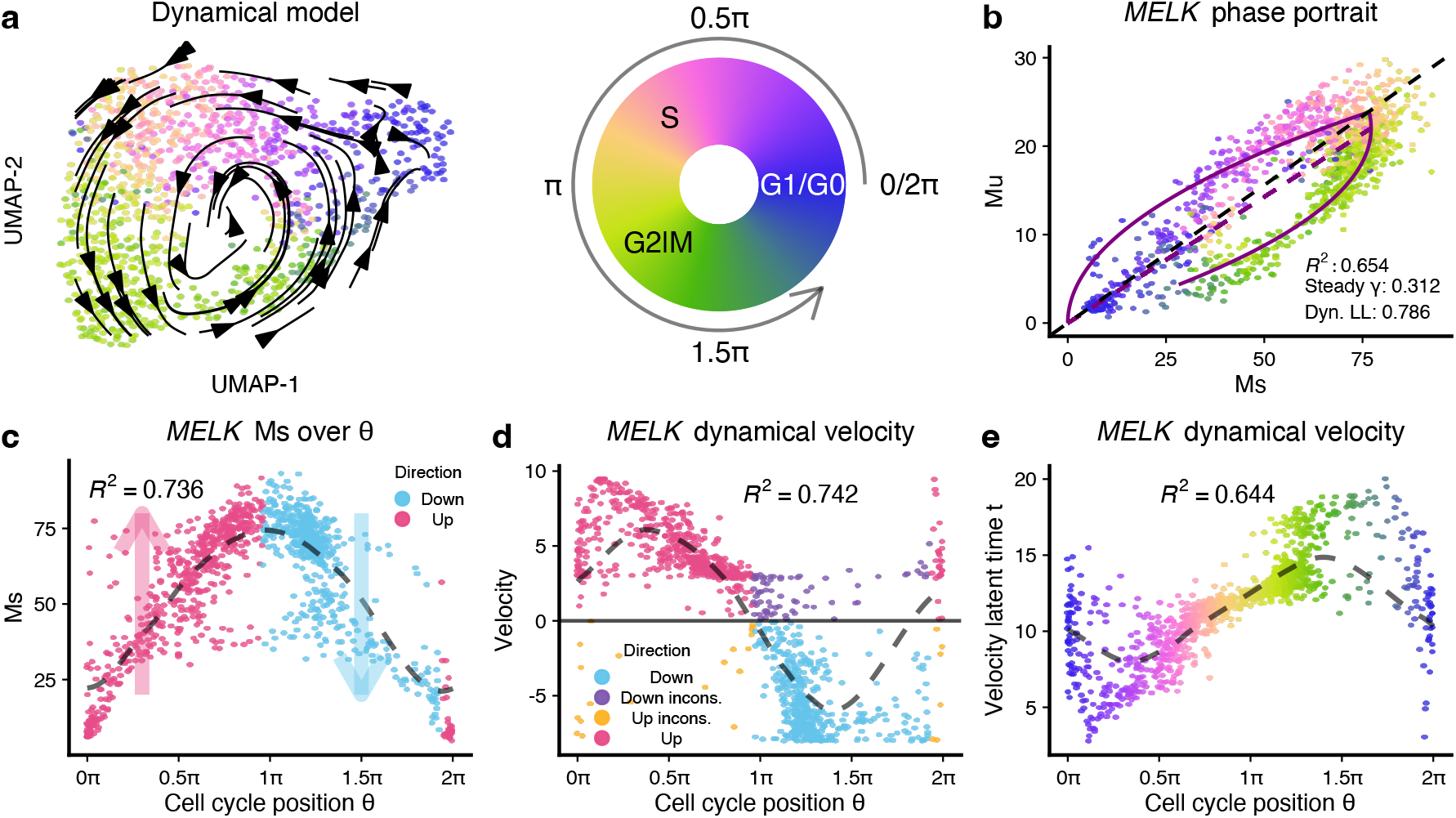
The RNA velocity application on FUCCI dataset. **(a)** RNA velocity vector field is visualized using the transition probability method on the UMAP embeddings of the FUCCI data. Each point represents a cell and is colored by the tricycle cell cycle position. **(b)** Phase portrait of gene *MELK*, of which the likelihood is the highest among all velocity genes inferred by the dynamical model. The purple lines represent the dynamics inferred by the dynamical model. **(c)** Scatter plot shows smoothed expression of *MELK* over cell cycle position. The dashed line is the fitted line by periodic loess (Methods). The expected direction of change is inferred on the fitted loess line and visualized by colors. **(d)** Scatter plot shows the estimated RNA velocity of *MELK* over cell cycle position. The signs of velocity estimations are compared to those inferred in (c), with inconsistent directions colored black. The variance explained by cell cycle position (*R*^2^ in the figure) is comparable for velocity values compared to Ms in (c). **(e)** The velocity latent time for *MELK* generally agrees with cell cycle position, except for cells around G1 or G0 phases.

Using our estimated cell cycle time from the tricycle, we can define the up- and down-regulation phases for each gene by smoothing expression using the cell cycle time loess fitted line (Figure 6c). Comparing these phases with the sign of the estimated velocities using the dynamical model reveals a high concordance (Figure 6d). A feature of the dynamical model is the estimate of a cell-specific latent time which has good concordance with cell cycle time for *MELK* (Figure 6e). Here, we use the gene-specific latent time, which is different from the gene-shared latent time(Bergen et al., 2020).

Using the steady-state model, we get essentially the same vector field (Supplementary Figure S12a). However, there are non-negligible inconsistencies between gene-level RNA velocity estimations from the steady-state and the dynamical model. For example, for the *MELK* gene 24% cells exhibit different directions of change (Supplementary Figure S12b-c) between the two models; the dynamical model agrees better with the direction of change inferred using cell cycle position. For each gene, we compute the PCC between the estimated velocities from the two models as well as the number of cells with inconsistent directions (Supplementary Figure S12d). Across genes, the median PCC between the models is only 40%, with 30% of cells showing an inconsistent direction of change. Intriguingly, the two models yield qualitatively similar vector fields despite the substantial differences in estimated velocities. This, again, raises the issue of to what extent the velocities actually impact the vector fields. We emphasize that this comparison only focuses on the direction of change and not the rate of change.

If we take all the cycle genes with a *R*^2^ ≥ 0.5 for Ms over the cell cycle position, we observe that the dynamical model is more consistent in the inference of velocity direction (Supplementary Figure S13).

Using the tricycle cell time can be criticized because it is inferred using the expression data. If we replace tricycle cell cycle time with FUCCI pseudo-time, we observe a higher degree of discordance with FUCCI pseudo-time (Supplementary Figure S14) compared to tricycle time. Supplementary Figure S15 shows 10 additional genes with similar behavior.

In summary, when it comes to the inference of direction of expression change, RNA velocity appears to work well in this cell cycle system, likely due to the fact that dynamic genes in this system experience both induction and decrease over the course of the cell cycle and potentially a better fitting to the underlying models of transcriptional regulation. We note the unique role of the cell cycle in velocity analysis: the periodic nature of the cell cycle fits well with the biophysical model of transcription, and cell cycle genes are often among the best fitting genes across biological systems. As a result, the velocity vector field is correct and the direction of change of the gene-level velocities is largely correct for the dynamical model. We find it noteworthy that the steady-state model also yields a correct velocity vector field despite the two models yielding high-dimensional velocity estimates with substantial inconsistencies. Based on this, we believe the cell cycle represents a system that is particularly well suited to RNA velocity, and we caution against generalizing its performance to other systems.

### A quality control measure for RNA velocity model

A central question is when to trust RNA velocity vector fields. To assess this, we focus on how well the estimated latent time explains variation in the expression estimates. In the dynamical model, we estimate a gene-specific latent time representing the unknown time parameter in the differential equations; this quantity is unavailable from the steadystate model. In a subsequent step, the gene-specific latent time is summarized into a gene-shared latent time by taking the quantile (across genes) of the estimated gene-specific latent times. We ask how much variation in expression is explained by a loess fit on the gene-specific latent time (Figure 7a,b). As an example, for one specific gene, for simulated data, we observe *R*^2^ = 0.73 using the estimated latent time compared to *R*^2^ = 0.52 using the true time. This suggests some degree of overfitting, possibly caused by either the preprocessing step or the model fit itself. Fortunately, in addition to the spliced counts, we also get unspliced counts, which are also smoothed but using the k-NN learned from the spliced counts. By comparing the difference between the spliced and unspliced expression matrices as a function of estimated latent time or the true time (Figure 7c,d compared to Figure 7a,b), we conclude that the source of the overfitting is the preprocessing step of smoothing using the spliced k-NN graph. We argue that this can be avoided by using the unspliced instead of the spliced matrix to assess how well RNA velocity model fitting works across genes because the preprocessing is done using the spliced k-NN graph and not the unspliced.

**Figure 7.**
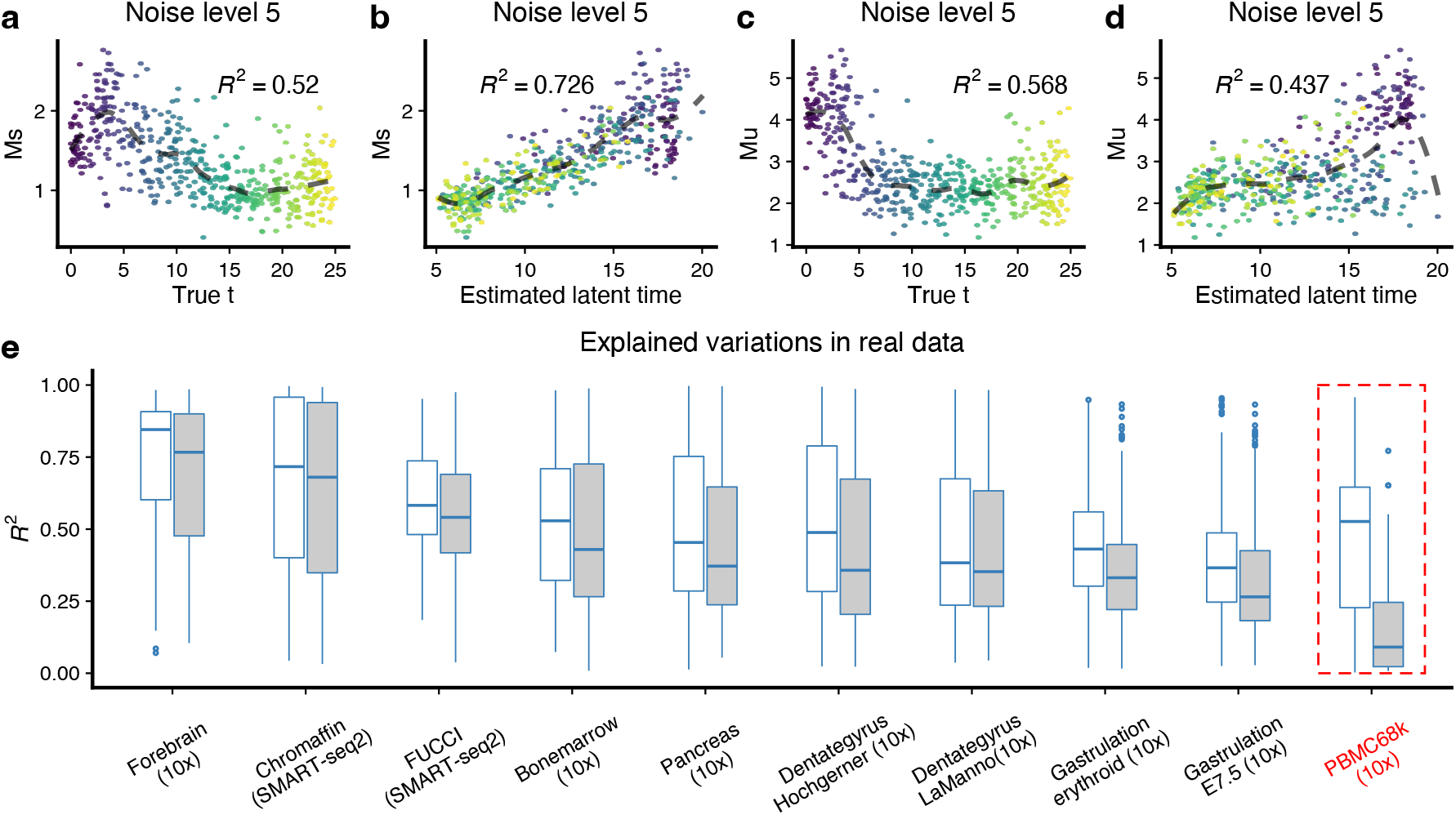
Using the *R*^2^ values to detect bad datasets with high noise. **(a-d)** Using a gene from the simulation at noise level 5, we show Ms over true latent time, Ms over estimated latent time, Mu over true latent time, and Mu over estimated latent time. The dashed lines are fitted loess, on which the *R*^2^ values are calculated (Methods). **(e)** *R*^2^ values of Ms and Mu over the estimated latent time for the top 300 velocity genes ranked by the likelihood in 10 real datasets. Note that in the PBMC68k dataset, where we know RNA velocity vector field does not work as we expect, the median *R*^2^ for Mu is less than 0.1, and the difference between the median *R*^2^ for Mu and Ms is about 0.5.

We applied this to the top 300 genes (ranked by the likelihoods of the dynamical model) in ten datasets (Figure 7e). The PBMC68k data – highlighted by Bergen et al. (2021) as a dataset where RNA velocity fails – is an outlier in this display, with a very high discrepancy between *R*^2^ for the spliced and unspliced expression values, and a very low *R*^2^ for the unspliced values. The two gastrulation data sets that have the second and third lowest *R*^2^ values have been reported to show unexpected directions in their RNA velocity vector fields that oppose the true differentiation path (Barile et al., 2021); the authors speculate that these inconsistencies are driven by time-dependent changes in expression dynamics. We interpret a small value of *R*^2^ for the unspliced matrix as evidence of a poor fit of the RNA velocity model. And we interpret a discrepancy between the *R*^2^ of the spliced and unspliced matrix to be evidence of overfitting arising from the k-NN smoothing. Altogether, this suggests that our measure has some functionality in the sense that extreme behavior on this measure should inspire low confidence in the vector fields.

## DISCUSSION

In physics, velocity is the combination of direction and speed and is defined as *v* = *ds*/*dt*, the derivative of position s with respect to time t. RNA velocity promises to bring temporal dynamics into gene expression analysis by estimating the derivative of the expression with respect to time. When assessing this promise, it is important to separately consider direction and speed (the length of the vector). For both direction and speed, RNA velocity produces two quantities: high-dimensional velocity vectors (each entry corresponds to a gene) and representations of these vectors in a suitable lowdimensional space, such as a UMAP embedding. To distinguish between these quantities, we refer to the low-dimensional representation as the “velocity vector field”, which is always relative to the choice of low-dimensional space and is easy to visualize.

### Why does RNA velocity appear to work?

The low-dimensional directions inferred by RNA velocity appear to be successful at describing known biology in many systems. “Describing known biology” is a qualitative statement reflecting that the visualized vector field reflects known progression through a system; an example is the pancreas dataset (Figure 2). We emphasize that this qualitative assessment is always exclusively in reference to the direction of the vector field; we will discuss speed below.

Most single-cell expression analyses – and all existing RNA velocity analyses – start by constructing a low dimensional embedding (usually) using UMAP. It is an open question under which conditions UMAP is ever guaranteed to reflect biological truth. Despite this lack of guarantees, it is indisputable that UMAP often achieves a representation that qualitatively reflects existing knowledge (related cell types and states are placed close together). We argue that any successful RNA velocity analysis starts with a UMAP which is deemed to represent existing or anticipated knowledge about the system. RNA velocity then produces a vector field overlay on this low-dimensional representation.

Here, we show that the direction of RNA velocity is strongly determined by the observed k-NN graph of the data. This k-NN graph is also directly reflected by the UMAP layout (indeed, one can view a UMAP layout as a representation of the k-NN graph). Together, this guarantees a *compatibility* between the UMAP and the vector field: vectors will always point towards neighbors on the UMAP. Importantly, this occurs regardless of whether the UMAP is in some sense “true”. Therefore, the directions inferred by RNA velocity cannot be interpreted as additional evidence for the correctness of the UMAP. For example, it is impossible for the RNA velocity vector field to construct a vector showing a transition between two cell populations that are distant on the UMAP. Instead, RNA velocity is more similar to a smoothing of the k-NN (and therefore UMAP) structure and cannot reveal a new structure but only depict a structure already present in the UMAP.

Our various experiments support these statements about the direction of the RNA velocity vector field; we next discuss these points and the experiments in detail.

### What influences the direction of RNA velocity?

First, we discuss the low-dimensional velocity vector field and focus on the particular case where mapping is done using velocity transition probabilities, as is common when visualizing RNA velocity on a UMAP, t-SNE, or other nonlinear embeddings. We show that the resulting vector field is strongly dependent on the structure of the k-NN graph (Figure 4). The use of the k-NN graph to preprocess (smooth) the spliced and unspliced matrix has a larger impact on the resulting vector field than the use of the k-NN graph to estimate velocity transition probabilities (Supplementary Figure S3). Additionally, we show how the resulting vectors are constrained to point exclusively toward sampled cell locations (Figure 2).

Second, we consider the direction of the lowdimensional RNA velocity vector field when the UMAP and k-NN graphs are perturbed away from the “true” structure. Our simulation results show that increasing noise can significantly perturb the observed (“learned”) k-NN graph away from the true neighborhood structure. This is best visualized by comparing the UMAP layouts at different noise levels (Supplementary Figure S2). While our observation is based on a specific simulation result, we hypothesize that this is a common phenomenon in single-cell expression data, where technical noise can be the greatest source of variation. In our simulations with moderate to high noise, we observe that a true one-dimensional manifold gets represented as a dense mass of cells on PCA or UMAP embeddings, where the quadrants of the mass are still broadly reflected specific time points, conditions, or cell states (Figure 4 and Supplementary Figure S2). Because specific cell states appear to occupy distinct regions of the dense mass of cells, and related cell types/states are placed close together, this is often interpreted as a signal in the data. Our simulations suggest that such a dense mass, caused by the noise of the data, might obscure a simple trajectory. In this scenario, RNA velocity produces a pleasing, smoothed, low-dimensional velocity vector field, which nevertheless is a wrong representation of the structure of the system (Figure 4). Given the importance of the observed k-NN graph in many methods in single-cell expression analysis, this observation may have far-reaching implications beyond RNA velocity analysis.

We now turn to the direction in high-dimensional space. First, we consider the relationship between high-dimensional velocities and the lowdimensional vector field. In our analysis of the FUCCI data, we show that both the dynamical model and the steady-state model yield overall similar vector fields, but that the inferred direction of change of expression is qualitatively different between the two models (Figure 6 and Supplementary Figure S12). This highlights that different high-dimensional velocities can result in the same low-dimensional vector field and cautions us against using the low-dimensional vector field to draw conclusions about the high-dimensional velocities. In our simulations, we observe that the estimated high-dimensional velocities strongly depend on the k-NN graph. Using the observed k-NN graph, there is the little concordance of directions between true high-dimensional velocities and the high-dimensional velocity estimates (Supplementary Figure S9). Together, this suggests that highdimensional velocities can be highly misleading. We note that high-dimensional velocities are not directly used in most RNA velocity analyses.

### RNA velocity does not estimate expression speed

We now turn our attention to expression speed: the length of the *v* = *ds*/*dt* vector. Speed is seldom directly analyzed, but precocious differentiation in disease may result from changes in speed. Again, there is a length of the high-dimensional vector and the length of the low-dimensional vector resulting from mapping into a low-dimensional embedding.

We show conclusively that there is a little-to-no relationship between high-dimensional vector length and length in the low-dimensional velocity vector field when velocities are mapped using transition probabilities (Figure 3). This suggests that vector length (speed) in the low-dimensional embedding is meaningless (in this scenario). This is explained by considering the velocity transition probabilities: high-dimensional vector length is simply not used to determine the low-dimensional vector length. The consistency between high- and lowdimensional vector lengths is substantially better when velocities are projected onto a PCA plot using an orthogonal projection operator. However, it is well appreciated that PCA plots in 2 dimensions regularly only capture part of the multiple biological processes co-occurring in single-cell data.

Our simulation experiments show that highdimensional velocity length estimates have poor concordance with true velocity vector lengths, although the concordance can be improved by using the true k-NN graph (Figure 5 and Supplementary Figure S8, S9). However, since the true k-NN graph is always unknown, it is of limited practical utility beyond the illustration of the discordance.

In summary, there is no reason to believe that current RNA velocity workflows are capable of estimating gene expression speed, neither in high nor in low dimensions. We make two comments related to speed. First, there is a significant gap in the ability to validate the speed of global gene expression transitions experimentally. If the accurate estimation of speed is of interest, it is critical to develop experimental approaches that will directly measure this property to provide a ground truth against which validation is possible. Second, there are issues with the theoretical definition of the speed of gene expression changes. When computing a vector length, coordinates (genes) contribute with equal weight. This makes sense in 3D physics but is less intuitively clear in expression analysis: for example, one might want to consider the expression level and intrinsic variation of each gene. In practice, we believe that a more useful concept of speed is one that is coupled to specific biological processes, such as cell cycle speed, differentiation speed, or transcriptional response to perturbation.

#### Limitations of our simulations

Following Bergen et al. (2021), we use a straightforward simulation strategy where all genes are velocity genes, and the dynamics fit the underlying ODE perfectly well. Furthermore, the gene-specific parameters are identical between the genes. This is highly unrealistic but provides an over-optimistic best-case scenario. Given the model failures in this simulation setup, we would expect even larger discrepancies with more real-world models. We have criticized – but not critically evaluated – the aggregation of genespecific latent times into a cell-specific latent time. Because the simulation setup imposes the exact ordering of cells for every gene, the aggregation step works well. To investigate issues with inferring a cell-specific latent time, we suggest one would need a simulation design where multiple processes are happening across multiple time scales, for example, cell cycling and differentiation happening at the same time.

#### The exception that proves the rule: the FUCCI data

Our analysis of the FUCCI data adds to the short list of attempts at validating RNA velocity using experimental data. We take advantage of cell cycle time (a useful time concept different from wall time). Using this approach, we find the relatively good performance of the dynamical model on cell cycle related genes in predicting the direction of gene expression changes (Supplementary Figure S13). Note that this is a partial validation: we are only considering the direction of change of expression (whether a gene is up- or down-regulated) and not whether the “speed”or the predicted new state is correct. There is a substantial discrepancy between the dynamical and the steady-state model, and both models arrive at the same vector field.

In our work, we have substantially criticized RNA velocity. Why do we maintain our criticism in light of our moderately successful validation attempt? We believe that the FUCCI data is a rare experimental system that fits the underlying RNA velocity model well. First, only a single underlying process is happening (cell cycle). This is a substantial simplification compared to many datasets where multiple processes occur simultaneously. Second, gene expression in this process follows a cyclic pattern with both up- and downregulation. This is in contrast to other processes, such as differentiation, in which most dynamic genes are either exclusively down- or up-regulated along the process. An example of such a system is red blood cell development, where the regulation of some key genes, such as hemoglobin genes, is monotonic (Sankaran and Orkin,2013). Interestingly, RNA velocity was recently reported to result in a reverse (wrong) direction in this system due to errors in assigning the correct expression phase (up/down) to key genes (Barile et al., 2021). Third, the UMAP suggests that the observed k-NN graph accurately reflects the underlying biology. Our criticism of RNA velocity is about the use of the method to “validate” a given embedding, and the FUCCI data does not really address this problem since we know the initial UMAP accurately reflects cell cycle progression.

### Summary

In light of our results, we believe that RNA velocity has far from achieving its stated goal: quantifying expression dynamics. Indeed, most applications of RNA velocity to date have exclusively relied on a qualitative interpretation of RNA velocity vector field estimates to ‘reinforce’ the validity of learned trajectories in a reduced dimensional embedding. We provide evidence here that this validation exercise is at best a circular logic and, at worst, potentially inaccurate and misleading. The promise of RNA velocity as a quantitative tool to examine expression dynamics further falls short when the validity of these estimates is explored. Speed is especially problematic and has received little attention in the literature. At its best, RNA velocity provides a potentially useful visualization tool, conceptually similar to pseudo-time ordering.

## METHODS

### Review of the dynamical model of RNA velocity for scRNA-seq data

RNA velocity is usually introduced through a pair of differential equations for the amount of spliced and unspliced RNA depending on the cell-specific latent time *t* for each gene independently (Bergen et al., 2020):

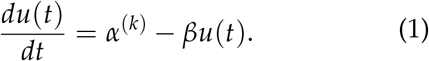

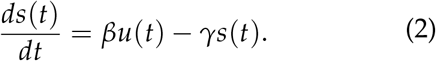

There are three unknown (constant) parameters for each gene: the transcription rate (*α*), the splicing rate (*β*), and the degradation rate (*γ*). Unlike the steady-state model, which only searches for the degradation rate *γ*, the dynamical model solves for the three parameters. The analytical solutions to equation 1 and equation 2, as given in Bergen et al. (2020), are the key parts for the estimation of the parameters:

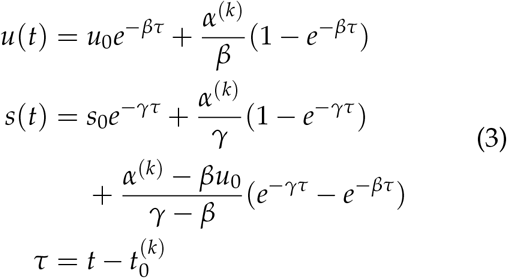

Here, *u*_0_ and *s*_0_ denote the initial unspliced and spliced counts, which are both set to 0 as default in implementation. And *τ* is the difference between the latent time *t* and the time point at which the phase change occurs. The greatest advancement of the dynamical model is that we get the cell-specific latent time *t* of each gene. Calculating the cellspecific velocity is performed by inserting the calculated values of *u*(*t*) and *s*(*t*) into the equation 2 instead of taking the residuals of the quantile regression as done by the steady-state model. We note that one of the important characteristics of the equation system 3 is that both *s*(*t*) and *u*(*t*) are univariate functions of cell-specific latent time *t* given all estimated parameters. It follows that equation 2 can also be expressed as an univariate function of the cell-specific latent time *t*.

### Real scRNA-seq datasets filtering and normalization

We got the Forebrain data (La Manno et al., 2018), the Bonemarrow data (Setty et al., 2019), the Dentategyrus LaManno data (Hochgerner et al., 2018; La Manno et al., 2018), the Pancreas data (Bastidas-Ponce et al., 2019; Bergen et al., 2020), the Gastrulation erythroid data (Pijuan-Sala et al., 2019), the Dentategyrus Hochgerner data (Hochgerner et al., 2018; Bergen et al., 2020), the Gastrulation E7.5 data (Pijuan-Sala et al., 2019), and the PBMC68k data (GXY Zheng et al., 2017) from the scVelo package directly (https://scvelo.readthedocs.io/api/#datasets). We got the Chromaffin data (Furlan et al., 2017; La Manno et al., 2018) from http://pklab.med.harvard.edu/velocyto/notebooks/R/chromaffin2.nb.html and the FUCCI data (Mahdessian et al., 2021) from https://drive.google.com/file/d/149ICTtieYjuKWZoLwRLzimwff0n6eWqw/view?usp=sharing. Usually, we did the following procedures for all real datasets: we first filtered genes with more than 20 counts across cells in both spliced and unspliced count matrices, and we only retained cells with more than 200 counts across genes in both spliced and unspliced count matrices; the filtered spliced count matrix was library size normalized across cells and log_2_ transformed by function normalizeCounts and then used as traditional expression matrix for PCA; we only used the top 2000 highly variable genes for PCA and the top 30 principal components for UMAP. All cell-type labels were included with the downloaded data.

For the following datasets, special treatments were applied. **FUCCI dataset**: For convenience, we scaled the FUCCI pseudotime to the range [0, 1] by dividing the pseudotime of each cell by the maximum pseudotime between cells. Cell cycle positions are estimated using the tricycle Bioconductor package (SC Zheng et al., 2022). **Dentategyrus LaManno and PBMC68k datasets:** Instead of using the sum of spliced and unspliced across cells threshold 20, we used threshold 30 as the number of cells is considerable for these two datasets.

### Construction of the k-NN graph

The k-NN graph is constructed using the pp.neighbors function in the scVelo package with all default parameters. Internally, it runs functions from the Python umap package, which searches for nearest neighbors in Euclidean distance of the top 30 PCs (default setting) and assigns weights to each edge (McInnes et al., 2018). (For simulations with the number of genes less than 500, the scVelo package will use the spliced counts matrix instead of PCs.) This results in an undirected weighted graph. We could use a symmetric *n* × *n* matrix **W**, of which column sums are normalized to 1, to represent such a k-NN graph.

### Smoothing of the count matrices

The smoothed count matrices Ms and Mu are calculated using the pp.moment function in the scVelo package with all default parameters. The smoothed count of a cell *i* for a particular gene *g* is given by the weighted sums of raw counts of k nearest neighbors:

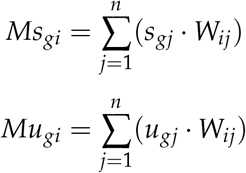

Note that the spliced and unspliced counts are smoothed independently, but we use the same weighted k-NN graph for both.

### Gene-level velocity estimation

We use the scVelo tl.velocity function with mode = ‘deterministic’ for steady-state model velocity estimation. For the dynamical model velocity estimation, we run the scVelo tl.recover_dynamics function to recover dynamics and then get the velocity values by the scVelo tl.velocity function with mode = ‘dynamical’. All parameters are default as in the scVelo package. In brief, the steady-state model fits an extreme quantile regression line and calculates the residuals for each gene, which are returned as the gene-level velocity values. Along with the velocity matrix, we also get the estimated degradation rate *γ* and the coefficient of determination for each gene. Only genes with a coefficient of determination greater than a pre-set threshold (0.01 used as default) will be labeled as velocity genes. Only these velocity genes will be used for visualization. Unlike the steady-state model, the dynamical model simultaneously solves the transcription rate *α*, the splicing rate *β*, and the degradation rate *γ*. The inference is also made for each gene independently. For each gene, the direct output of the dynamical model is the gene parameters and a vector of latent time *t*, whose length is the number of cells n. The gene-level velocity value is then calculated using the system of equations 3 and the equation 2. Note that we can only recover the parameters for a subset of genes, within which passes the pre-set threshold of coefficient of determination and likelihood are labeled as the velocity genes.

### Simulation settings

The simulation strategy is the same as described in Bergen et al. (2021), as we used the simulation function in the scVelo package. Briefly, we simulate the spliced count and unspliced count matrices based on the system of equations 3. Note that the simulation process is independent for each gene, and it is trivial to get *s*(*t*) and *u*(*t*) as long as we have assigned other parameters. We use the transcription rate *α* = 5, the splicing rate *β* = 0.3, and the degradation rate *γ* = 0.5 for all genes, as used in Bergen et al. (2021). To simulate data with a number of cells *n*, a cell-specific pseudotime vector is generated log uniformly distributed and scaled to the range [0, *t_max_*], with *t_max_* always set to 25 in our simulations. To make the counts different across genes, we further rescale the cell-specific time vector *t* to some interval within [0, *t_max_*]. This step will keep the orders of cells, but make all cells only cover some part of the full dynamics for a given gene. The same procedure is repeated for each gene to obtain a latent time matrix with *m* rows (*m* genes) and *n* columns (*n* cells). We can then plug the gene latent time matrix into the system of equations 3 to obtain the spliced count and unspliced count matrices. The true RNA velocity matrix comes naturally from the equation 2. After the generation of spliced and unspliced matrices, we add Gaussian noise with mean 0 and standard deviation *σ* to the theoretical spliced and unspliced matrices to make the simulated data used in our analyses. The standard deviation *σ* is equal to the noise level multiplied by the 99% percentile of the spliced or unspliced counts divided by 10 in our manuscript and in Bergen et al. (2021). Note that, as in Bergen et al. (2021), we do not perform library size normalization and log_2_ transformation on the spliced count matrix. The k-NN graph construction and smoothing of counts matrices are performed as described previously. In the simulation where we use another “true” k-NN graph, the k-NN is calculated before we add Gaussian noise to the raw spliced and unspliced counts. Both the steady-state model RNA velocity and the dynamical model RNA velocity are inferred using the scVelo package, resulting in a velocity matrix *V* to be used later. Also, we force the labels of the “velocity gene” of all genes to be true, as in Bergen et al. (2021). For the steady-state model, the estimated velocity of all *m* genes will be used for later visualization. For the dynamical model, some of the genes are still excluded for visualization due to missing velocity estimations from the dynamical model. Coercion could potentially improve visualization results since we know that all genes are true “velocity genes”.

### Mapping high-dimensional velocities into a low-dimensional embedding

After getting the velocity matrix, which contains a velocity value for *m*^*^ (*m*^*^ is the “velocity gene” that passes predefined thresholds) genes and *n* cells, we need to map the high-dimensional velocity into the same low-dimensional embedding of expression (spliced counts). In our manuscript, we use the following three methods to map high-dimensional velocities, with the first two used by La Manno et al. (2018). We use **S** to represent the expression matrix (raw spliced counts matrix), which has the shape of m rows (genes) and n columns (cells). Note that we describe the precise procedures here, which might look slightly different from the simplified version in the Results section.

#### Direct projection in PCA

We have used this method only on simulated data with known true velocity. Theoretically, this method could be used in any embedding methods with a linear operator 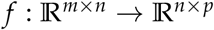. Specifically for PCA, we have

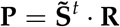

where **R** represents the m-by-p rotation matrix; 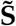 is a m-by-n spliced count matrix with row-means centered. The resulting n-by-p **P** is the cell-level principal components matrix. We can map velocities as

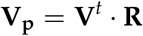

The formulation here is much simpler than the application to real data, as we omit library size normalization, log_2_ transformation, and highly variable gene selection. Note that in the simple simulation setting and PCA space, we project the velocity vectors directly, which is (almost) equivalent to taking the difference between the future state (**S**^*^denotes the future expression state matrix) and current state:

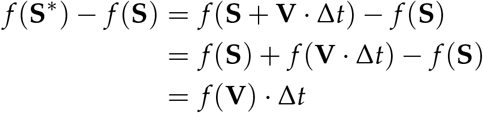

*f* (**S**^*^) – *f* (**S**) = *f*(**V**) when Δ*t* = 1. It is clear that Δ*t* is a trivial scalar, since all vectors are rescaled in the final visualization. However, the choice of Δ*t* would matter in real data as non-linear operations are involved, such as normalization of the library size and transformation of log_2_.

#### Velocity transition probability method

The velocity transition probability method was first introduced by La Manno et al. (2018) and was reused by Bergen et al. (2020) with some modifications. We use the implementation by Bergen et al. (2020) with all default parameters in the scVelo package.

For a given cell *i*, we have its velocity vector *V_i_* and its expression vector *Ms_i_* (the length for both two is *m*^*^ since we only consider the “velocity” genes filtered by scVelo). Also, note that the Ms matrix is being used for the transition probability instead of the raw spliced count matrix. The first step to get the velocity transition probability matrix is the calculation of a n-by-n cosine similarity matrix. The advantage of using cosine similarity or PCC to quantify the relationship between cells is that we do not need to choose Δ*t*. For a cell *i*, we consider cells which are cell *i*’s k-nearest neighbors and recursive k-nearest neighbors, denoted as {*n*(*n*(*i,k*), *k*)}. The k-NN graph is precalculated on the PCA of the raw spliced count matrix and was used for Ms and Mu. The cosine similarity between cell *i* and cell *j* is given as

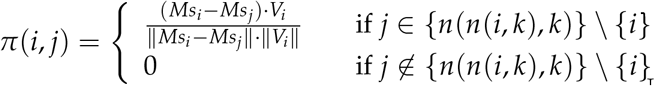

The exponential kernel is then applied on the cosine similarity matrix to get the velocity transition matrix. Specifically, the transition probability from cell *i* to cell *j* is given as

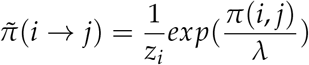

with *z_i_* as the cell normalization factor 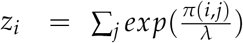 and the constant kernel width parameter *λ*. There are optional variance stabilization transformations mentioned in La Manno et al. (2018) and Bergen et al. (2020), but we use the default parameter in scVelo, which does not perform any variance stabilization transformations. Given a embedding **Q**, the normalized location difference between cell *i* and *j* is 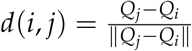. The mapped low-dimensional vector for cell *i* is calculated as

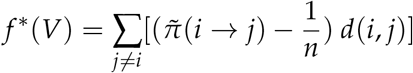

where *f*^*^ is a non-existing symbolic function. The idea behind the velocity transition probability is to weigh the likelihood that the cell *i* will become the cell *j* in the future state.

#### UMAP transform method

The UMAP transform method is an experimental method that has not been used in previous papers on RNA velocity (La Manno et al., 2018; Bergen et al., 2020). Unfortunately, we were unable to systematically confirm the correctness of the method since the UMAP transform function itself has not been systematically evaluated. To illustrate, we use *f* to represent the function that projects data into the PCA space. Since we use the PCA results as the input of UMAP function, the UMAP embedding is given as *g*(*f*(**S**)) = **Q**. Note that we use the function 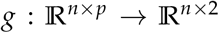 to represent the UMAP embedding process, but g is not as straightforward as the function f. Before we transform new points into the existing UMAP space, we need to get new PCA coordinates, which requires us to choose a Δ*t*. The coordinates of the future states in PCA are given by *f*(**S**\*) = *f*(**S**+**V**·Δ*t*) = *f*(**S**) + *f*(**V**·Δ*t*). Thus, the coordinates of future states in UMAP are given as

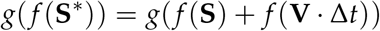

Note that *g* is not a linear function, so we could not expand the right part. Finally, we map the highdimensional velocities into the low-dimensional UMAP space as

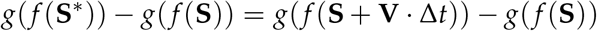

In our analysis, we always use Δ*t* = 1. We admit that this is a somewhat random choice, but it is hard to argue the choice because the fuzzy definition and poor interpretability of pseudotime/latent time t inherits from the RNA velocity models.

### Vector field visualization

After getting the low-dimensional cell-level vector field, we need to process it further as we could not show that many vectors (arrows) due to overplotting. We use two visualization strategies for the final visualization: the gridding method and streamline plot. While the streamline plot is aesthetically more appealing, we turn to the gridding method whenever we want to highlight more details. Again, as we have mentioned previously, the same vector field may look quite different when comparing the two methods. For all vector fields in the manuscript, we adapt the streamline plot and the gridding method implemented in the velociraptor R package (Rue-Albrecht et al., 2021) (Supplementary Figure S17 is an exception). The streamline plot connects vectors flowing towards and into a similar direction. For the streamline plot, which is used for real datasets, we use a resolution between 13 to 20. The gridding method takes the average of all vectors in the grid box. The averaged vectors are further scaled to look good based on the axis range. For the gridding method, we use a resolution of 20 for simulated data and 30 for the pancreas data. We note that there is no existing metric to guide us in choosing the best resolution, so we have to choose a resolution that, we think, makes sense based on the number of data points and the embedding structure to balance the details and overplotting.

### Calculation of the vector length (speed)

The vector length (speed) in both high-dimensional space and low-dimensional space is defined as the *ℓ*-2 norm of the vector of each cell. Specifically, for high-dimensional velocities, the vector length of cell *i* is 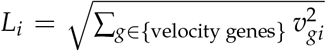 with *v_gi_* the velocity estimation of gene *g*. For low-dimensional embedding, the vector length of cell *i* is 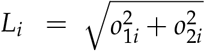 with *o*_1_*i* the mapped vector in the first dimension and *o*_2_*i* the mapped vector in the second dimension.

### Calculation of explained variation *R*^2^ of the fitted loess model

The calculation of the coefficient of determination *R*^2^ of the fitted loess model is given by

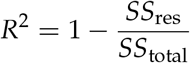

Here 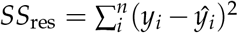 and 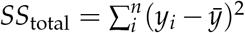. For the FUCCI pseudotime or tricycle cell cycle position, as we they track a full cell cycle, we fit a periodic loess model *y* ~ *t*, with *y* as any response variable, such as the Ms, or Mu, by concatenating triple *y* and triple FUCCI pseudotime *t* with one period shift to form [*y, y, y*] and [*t* – 1, *t, t* + 1] (or [*t* – 2*π, t, t* + 2*π*] for cell cycle position). Note that instead of using all three copies of data points, we restrict the calculation of *SS*_res_ and *SS*_total_ on the original data points (the middle copy). For the velocity latent time, we could not decide whether the latent time spans a complete period. Thus, we fit normal loess model for the velocity latent time.

### Calculation of Normalized Root Mean Square Error (NRMSE)

In the simulations, we know the true values, such as the true RNA velocity at the gene level. We use both PCC and NRMSE to quantify how good the estimations are. The NRMSE is calculated as

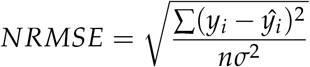

Here, *y_i_* represents the true values; *ŷ_i_* are the estimated values; *n* is the number of observations. The standard deviation of the true values *σ* is used to normalize the root mean square error for making different simulations comparable.

### Data and code availability

The Forebrain data (La Manno et al., 2018), the Bonemarrow data (Setty et al., 2019), the Dentategyrus LaManno data (Hochgerner et al., 2018; La Manno et al., 2018), the Pancreas data (Bastidas-Ponce et al., 2019; Bergen et al., 2020), the Gastrulation erythroid data (Pijuan-Sala et al., 2019), the Dentategyrus Hochgerner data (Hochgerner et al., 2018; Bergen et al., 2020), the Gastrulation E7.5 data (Pijuan-Sala et al., 2019), and the PBMC68k data (GXY Zheng et al., 2017) are all available from the scVelo package directly at https://scvelo.readthedocs.io/api/#datasets (Bergen et al., 2020). The Chromaffin data (Furlan et al., 2017; La Manno et al., 2018) is available at http://pklab.med.harvard.edu/velocyto/notebooks/R/chromaffin2.nb.html.

The FUCCI data (Mahdessian et al., 2021) at https://drive.google.com/file/d/149ICTtieYjuKWZoLwRLzimwff0n6eWqw/view?usp=sharing.

All the code to analyze the data and generate figures is available at https://github.com/hansenlab/RNAVelocityCode (SC Zheng, 2022).

## Disclaimer

The content is solely the authors’ responsibility and does not necessarily represent the official views of the National Institutes of Health or the National Science Foundation.

## SUPPLEMENTARY MATERIALS

### 1 SUPPLEMENTARY FIGURES

**Supplementary Figure S1.**
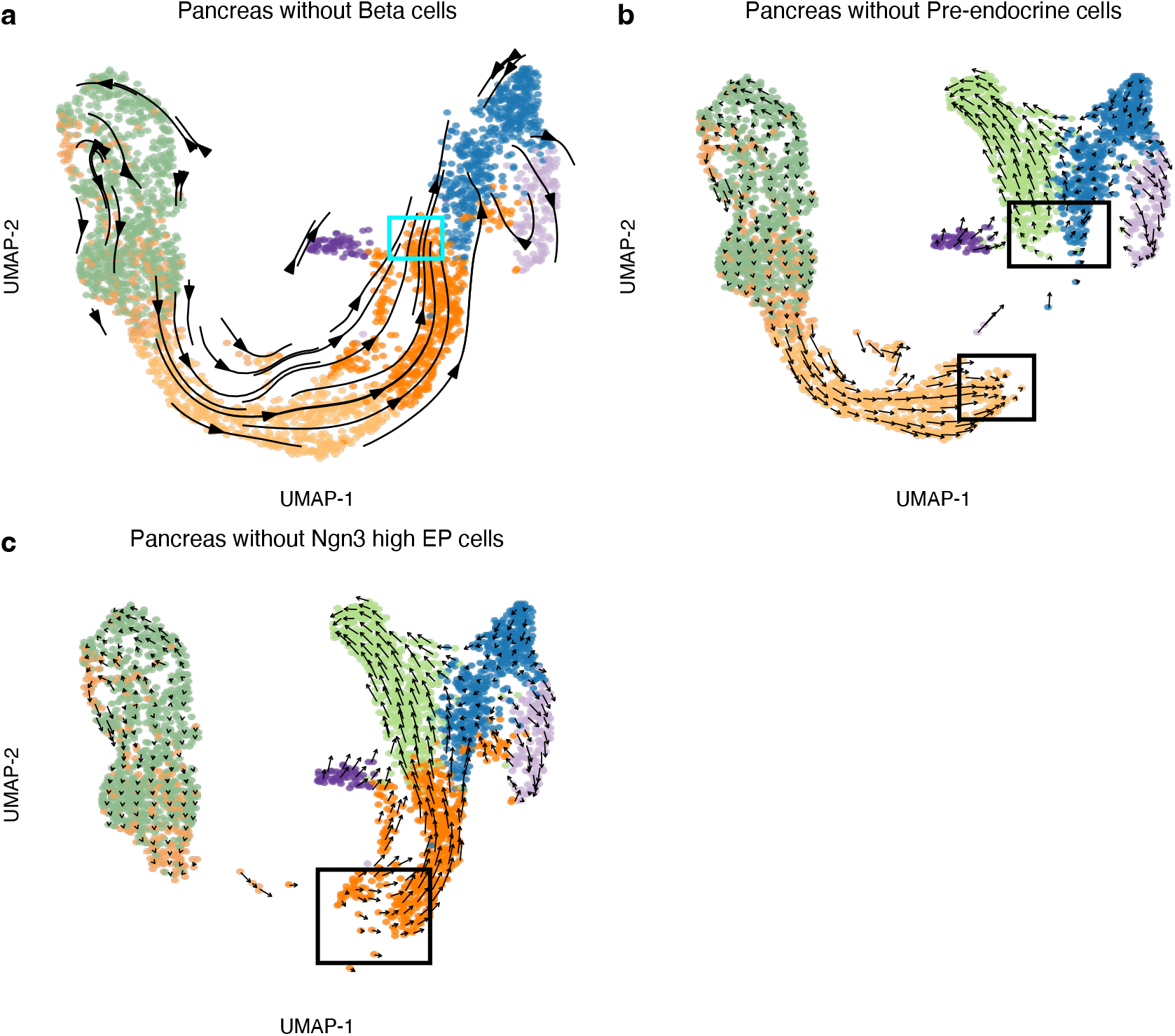
Embedded RNA velocity in the pancreas dataset after removing a cell type. **(a)** Dynamical model based RNA velocity is visualized on the UMAP for the pancreas data after removing the beta cells. **(b)** As (a), but the pre-endocrine cells are removed instead. **(c)** As (a), but the Ngn3 high EP cells are removed instead. Note that the arrows disappear for cells in the black rectangles of (b) and (c).

**Supplementary Figure S2.**
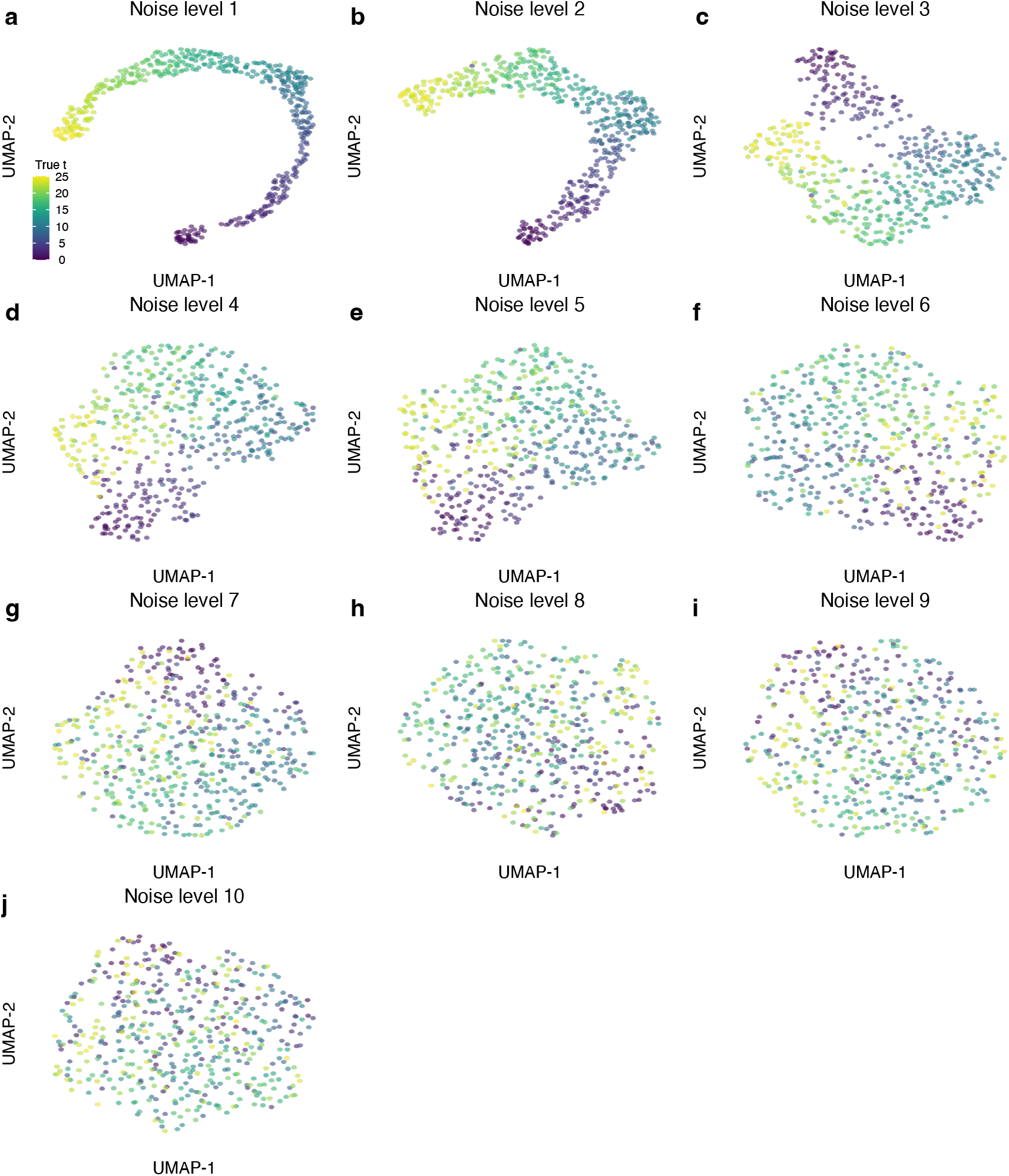
UMAP representation of the simulated data with different noise levels. Each point represents a cell, colored by the true latent time. We use a different noise level in each panel, increasing from 1 to 10. Similar to the PCA representation, the UMAP representation becomes like a big blob more and more with the noise level increases.

**Supplementary Figure S3.**
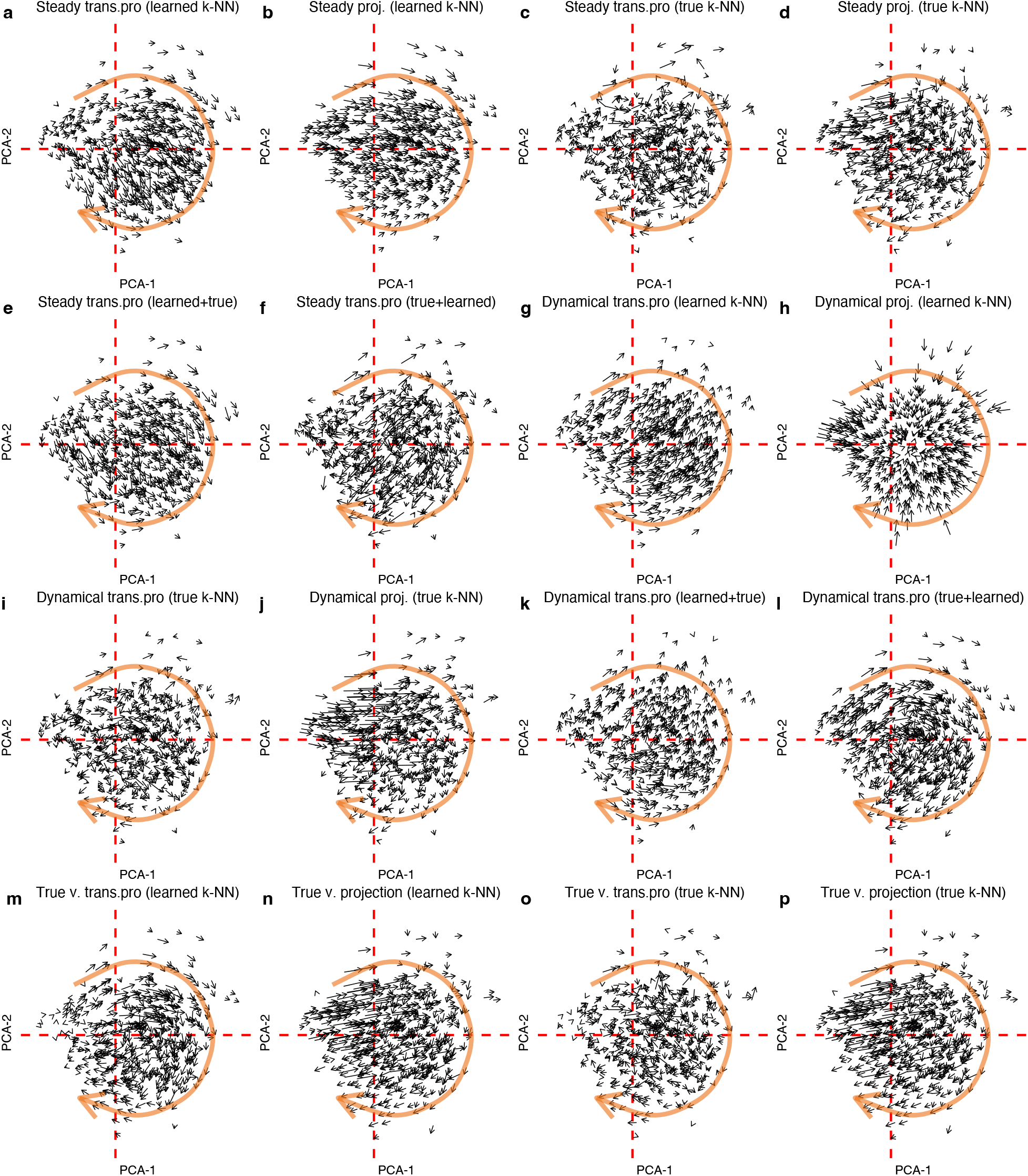
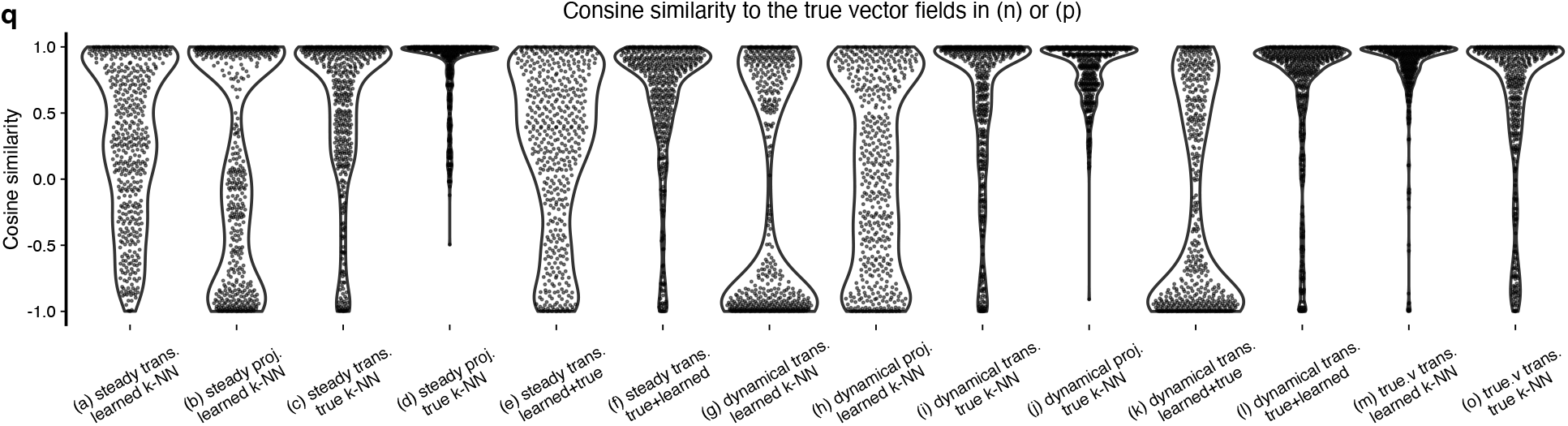
Visualized RNA velocity vector field using simulated data at noise level 5. **(a-p)** Visualized vector fields in all combinations of the following: types of high-dimensional velocities (estimated high-dimensional velocities from the steady-state model or estimated high-dimensional velocities from the dynamical model, or true high-dimensional velocities), mapping methods (transition probability or direct PCA projection), types of k-NN used for preprocessing (learned k-NN or true k-NN), and types of k-NN used for calculation of transition probability (learned k-NN or true k-NN). In (a-p), the type of velocities is given first in the panel title and followed by the mapping methods (trans.pro for transition probability and proj. for direct PCA projection). The type of k-NN is given in the parenthesis of each panel title. If there is one type of k-NN, then that type of k-NN is used for smoothing and transition probability calculation. If two types of k-NN are given, the first k-NN is used for smoothing and the second for transition probability calculation. In all panels, each point represents a cell. The big orange arrow approximates the true direction of the trajectory. Note that (n) and (p) is the same because the true velocity is unrelated to smoothing. **(q)** The cosine similarities between the mapped cell-level vectors and the “true” mapped cell-level vectors in (n) or (p).

**Supplementary Figure S4.**
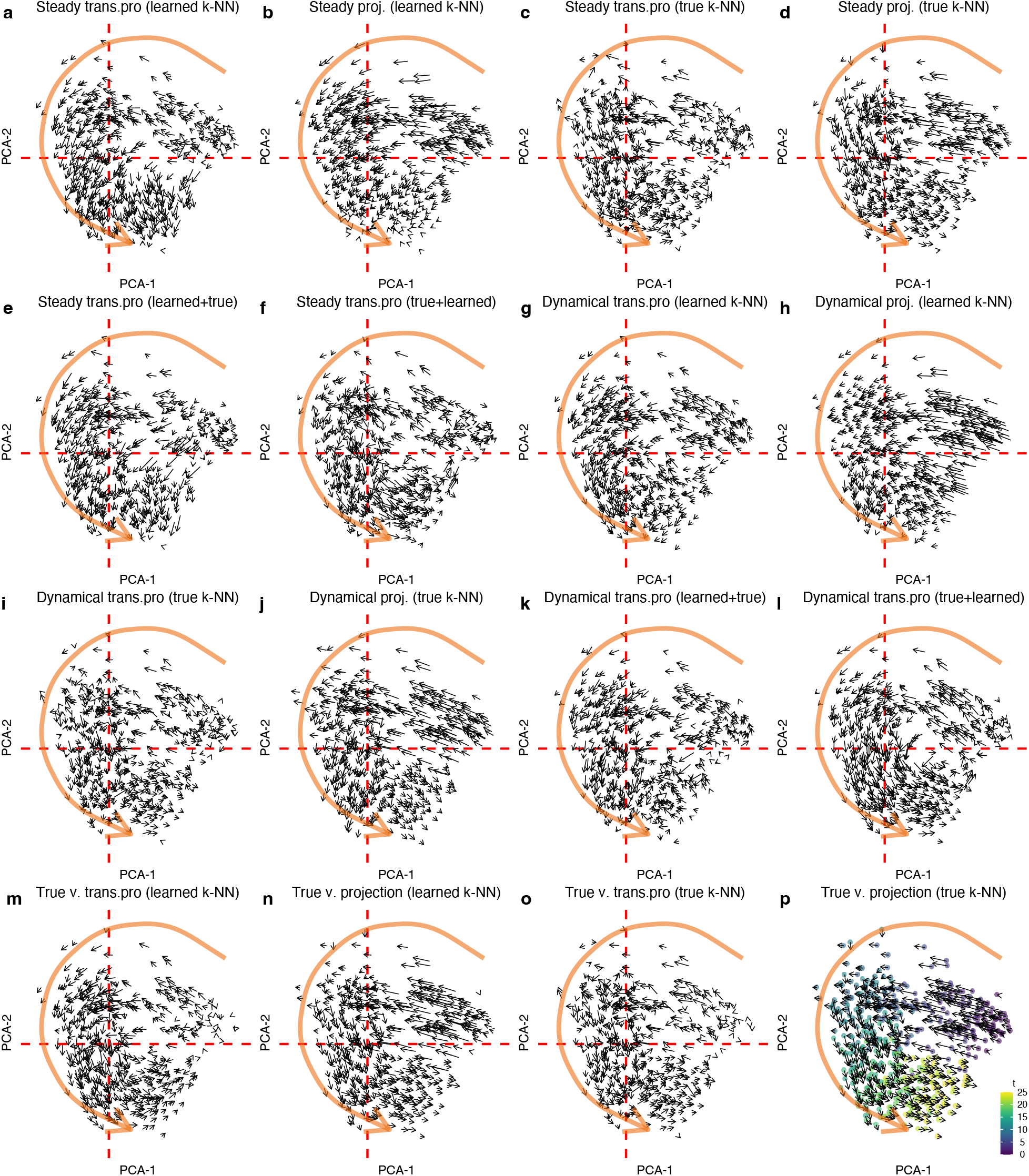
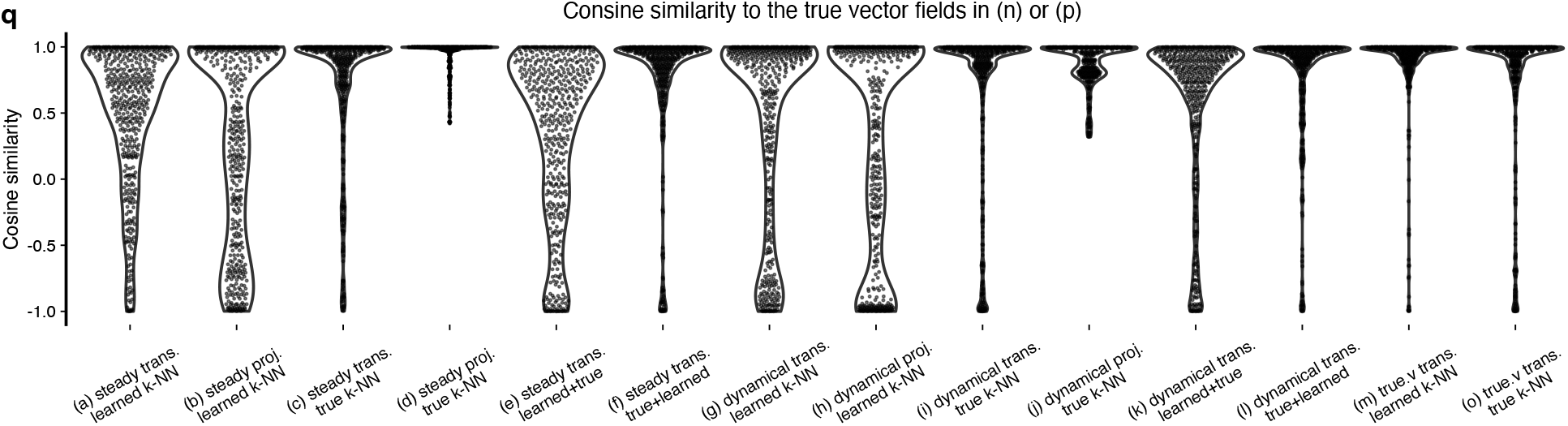
Visualized RNA velocity vector fields using simulated data at noise level 3. Similar to Supplementary Figure S3, but the noise level is lowered to 3. **(a-p)** Visualized vector fields in all combinations of the following: types of high-dimensional velocities (estimated high-dimensional velocities from the steady-state model or estimated high-dimensional velocities from the dynamical model, or true high-dimensional velocities), mapping methods (transition probability or direct PCA projection), types of k-NN used for preprocessing (learned k-NN or true k-NN), and types of k-NN used for calculation of transition probability (learned k-NN or true k-NN). In (a-p), the type of velocities is given first in the panel title and followed by the mapping methods (trans.pro for transition probability and proj. for direct PCA projection). The type of k-NN is given in the parenthesis of each panel title. If there is one type of k-NN, then that type of k-NN is used for smoothing and transition probability calculation. If two types of k-NN are given, the first k-NN is used for smoothing and the second for transition probability calculation. In all panels, each point represents a cell. The big orange arrow approximates the true direction of the trajectory. Note that (n) and (p) is the same because the true velocity is unrelated to smoothing. **(q)** The cosine similarities between the mapped cell-level vectors and the “true” mapped cell-level vectors in (n) or (p).

**Supplementary Figure S5.**
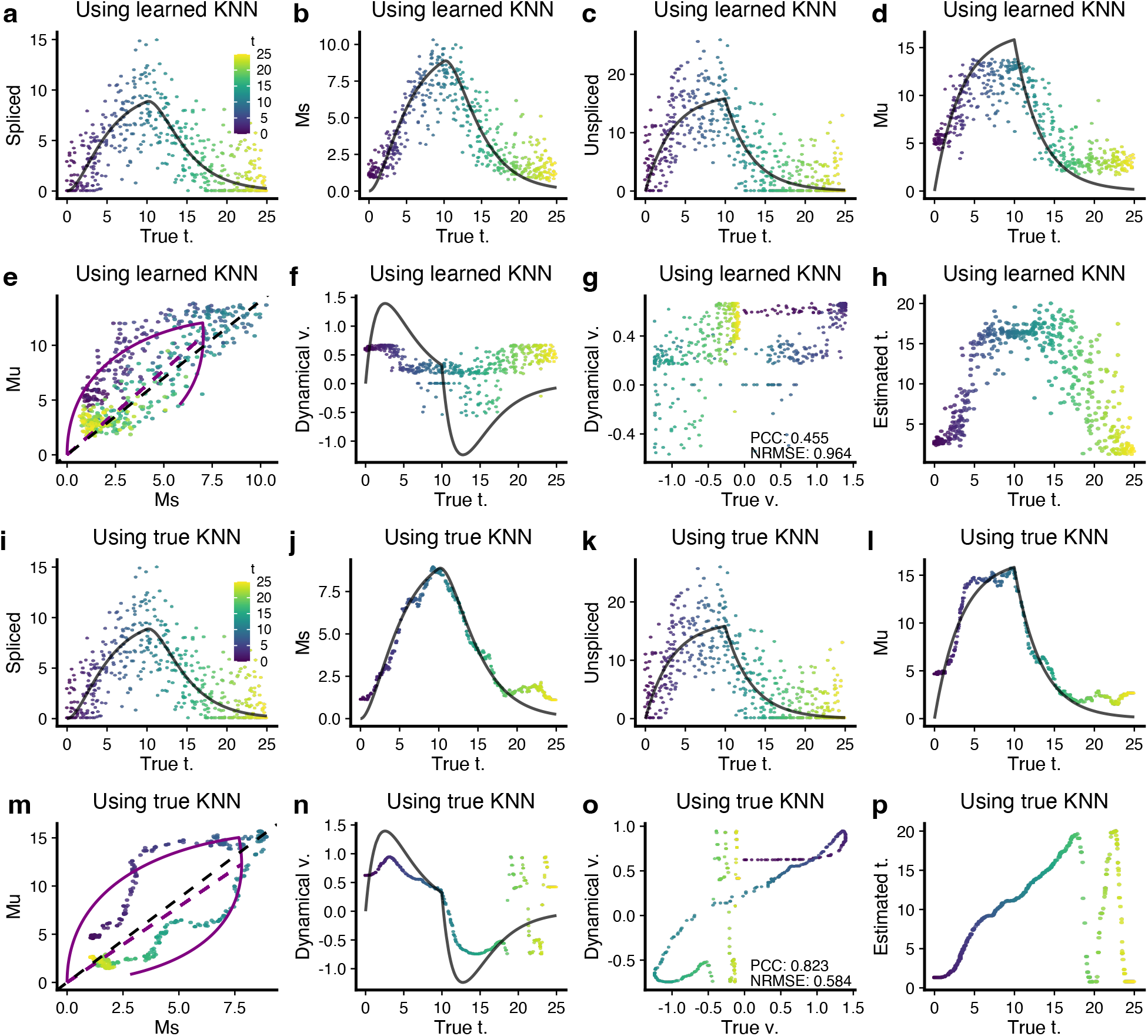
Gene-level RNA velocity estimation analyses using simulated data at noise level 3. This figure complements Figure 5. In all panels, a data point represents a cell and is colored by the known true latent time *t*. All solid black lines represent the known true values. **(a-d)** Scatter plots show the spliced counts, Ms, unspliced counts, and Mu over the true latent time *t* for a random gene in simulation. **(e)** Phase portrait shows the Ms over Mu for the same gene. The dynamics are estimated by the dynamical model. **(f)** Estimated velocity (points) and true velocity (black line) over true latent time *t*. **(g)** Scatter plot compares the estimated velocity values to the true velocity values. PCC and Normalized Root Mean Square Error (NRMSE) are given. **(h)** Scatter plot compares the estimated latent time to the true latent time. **(i-p)** As (a-h), but now we use the true k-NN to get Ms and Mu matrices. The estimated velocity values are much closer to the true velocity values with PCC 0.823 and NRMSE 0.584. Note that (a) and (i) are identical, and (c) and (k) are identical, too, since the k-NN does not affect raw counts.

**Supplementary Figure S6.**
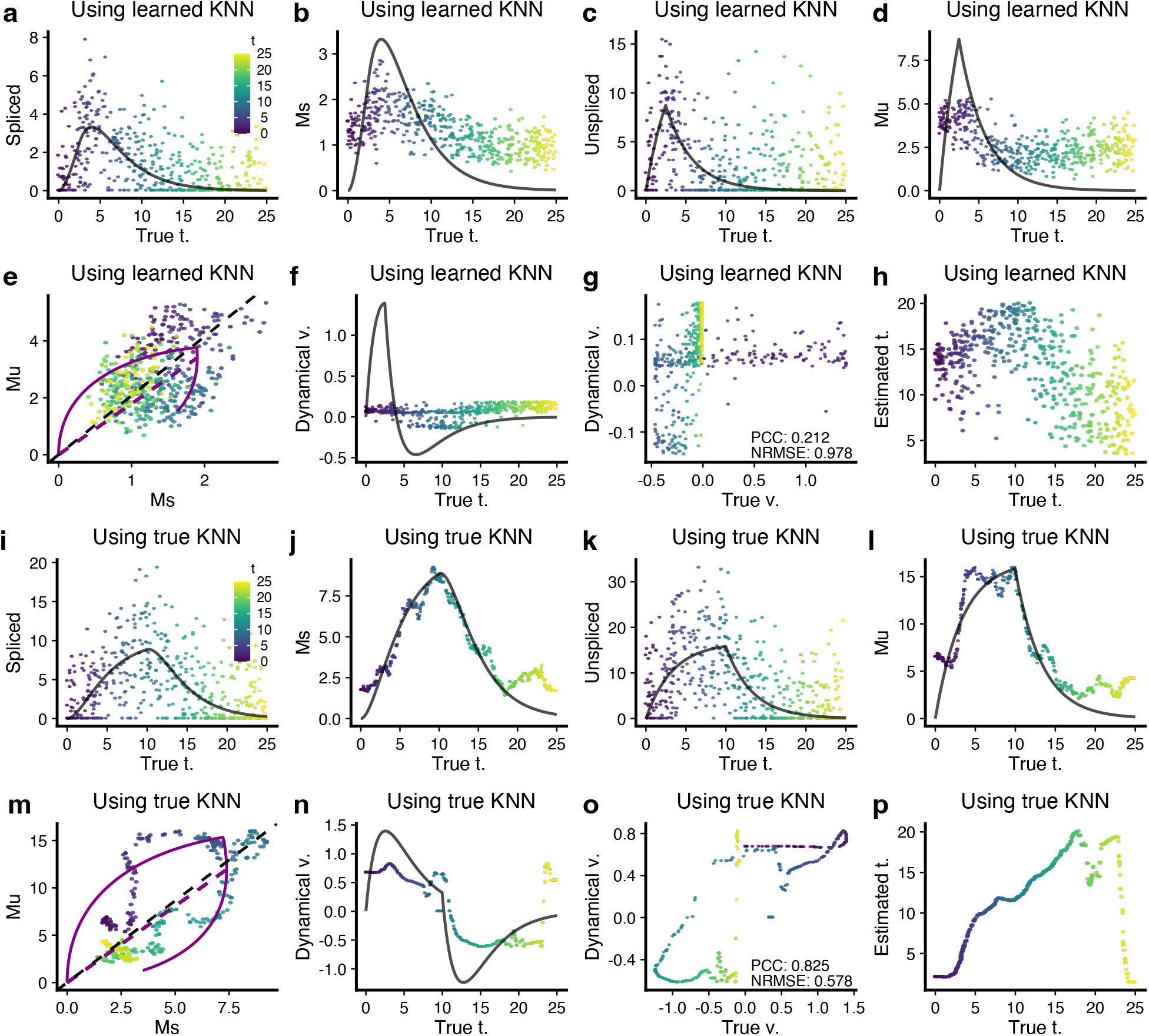
Gene-level RNA velocity estimation analyses using simulated data at noise level 5. This figure is similar to Supplementary Figure S5, but the noise level is 5. In all panels, a data point represents a cell and is colored by the known true latent time *t*. All solid black lines represent the known true values. **(a-d)** Scatter plots show the spliced counts, Ms, unspliced counts, and Mu over the true latent time *t* for a random gene in simulation. **(e)** Phase portrait shows the Ms over Mu for the same gene. The dynamics are estimated by the dynamical model. **(f)** Estimated velocity (points) and true velocity (black line) over true latent time *t*. **(g)** Scatter plot compares the estimated velocity values to the true velocity values. PCC and Normalized Root Mean Square Error (NRMSE) are given. **(h)** Scatter plot compares the estimated latent time to the true latent time. **(i-p)** As (a-h), but now we use the true k-NN to get Ms and Mu matrices. The estimated velocity values are much closer to the true velocity values with PCC 0.825 and NRMSE 0.578

**Supplementary Figure S7.**
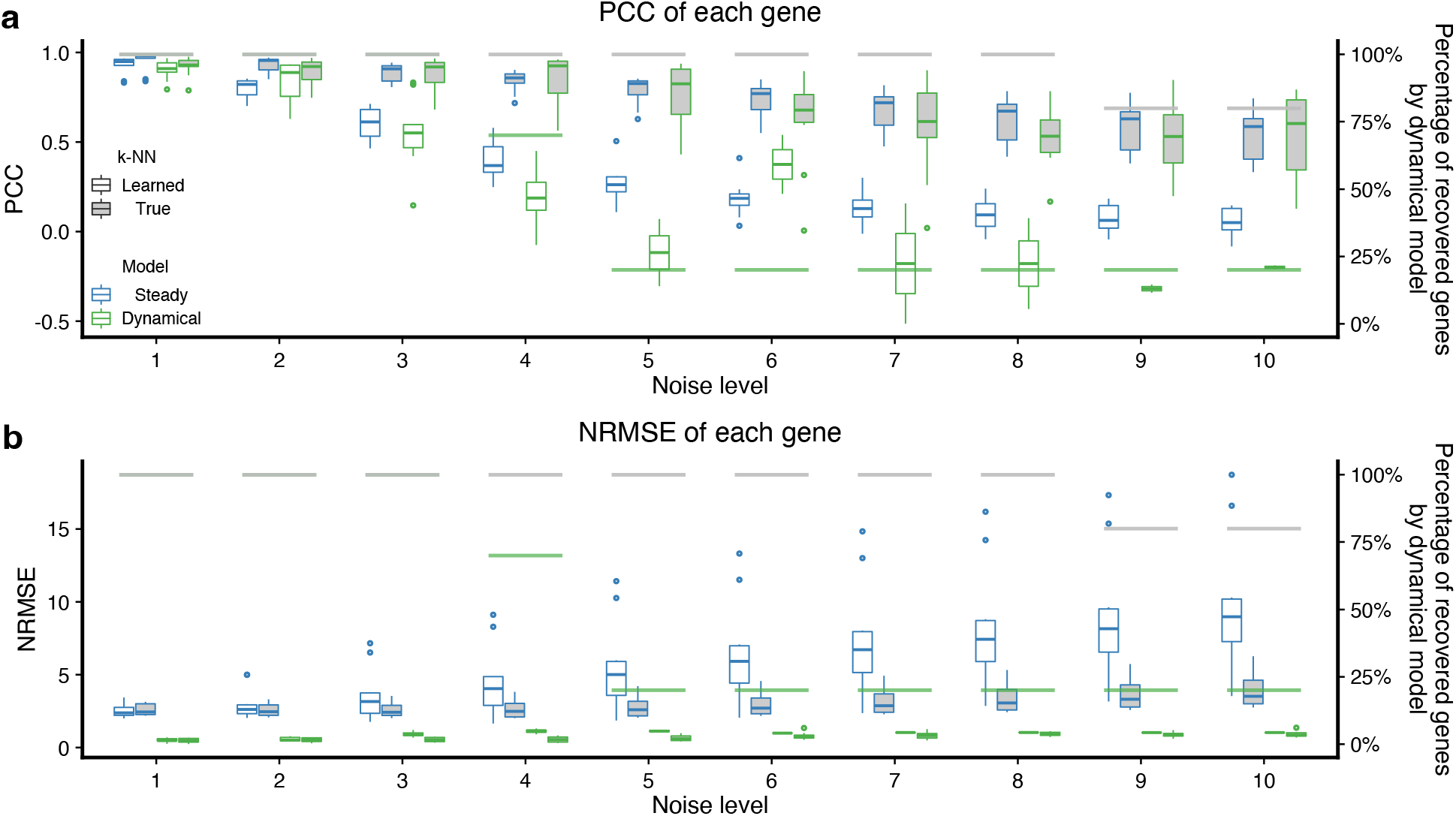
Comprehensive gene-by-gene evaluations of high-dimensional velocity estimations using simulations. **(a)** Boxplots show PCC between the estimated velocities and the true velocities of each gene. **(b)** Boxplots show NRMSE between the estimated velocities and the true velocities of each gene. All boxes show the left *y*-axis values and indicate the 25th and 75th percentiles. Whiskers extend to the largest values no further than 1.5 × interquartile range (IQR) from these percentiles. The grey and green horizontal lines correspond to the percentage of recovered genes by the dynamical model using the true and learned k-NN graph, respectively.

**Supplementary Figure S8.**
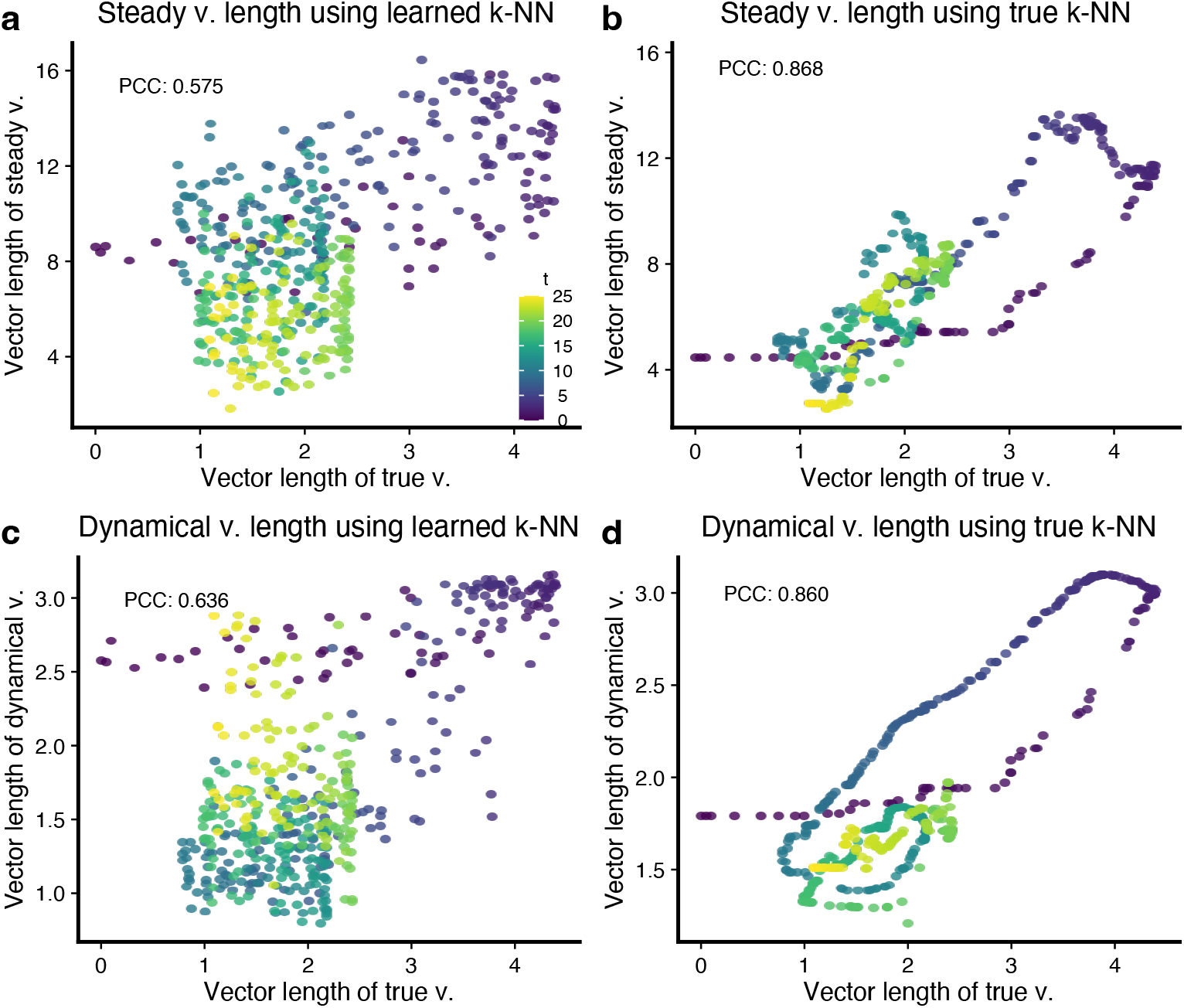
Comparisons of high dimensional velocity vector length at noise level 3. **(a)** Scatter plot compares the vector length of true (high-dimensional) velocity to that of the estimated (high-dimensional) velocity by the steady-state model using the learned k-NN graph. **(b)** As (a), but we use the true k-NN to infer (high-dimensional) velocity. **(c-d)** As (a-b), but the velocities are estimated using the dynamical model.

**Supplementary Figure S9.**
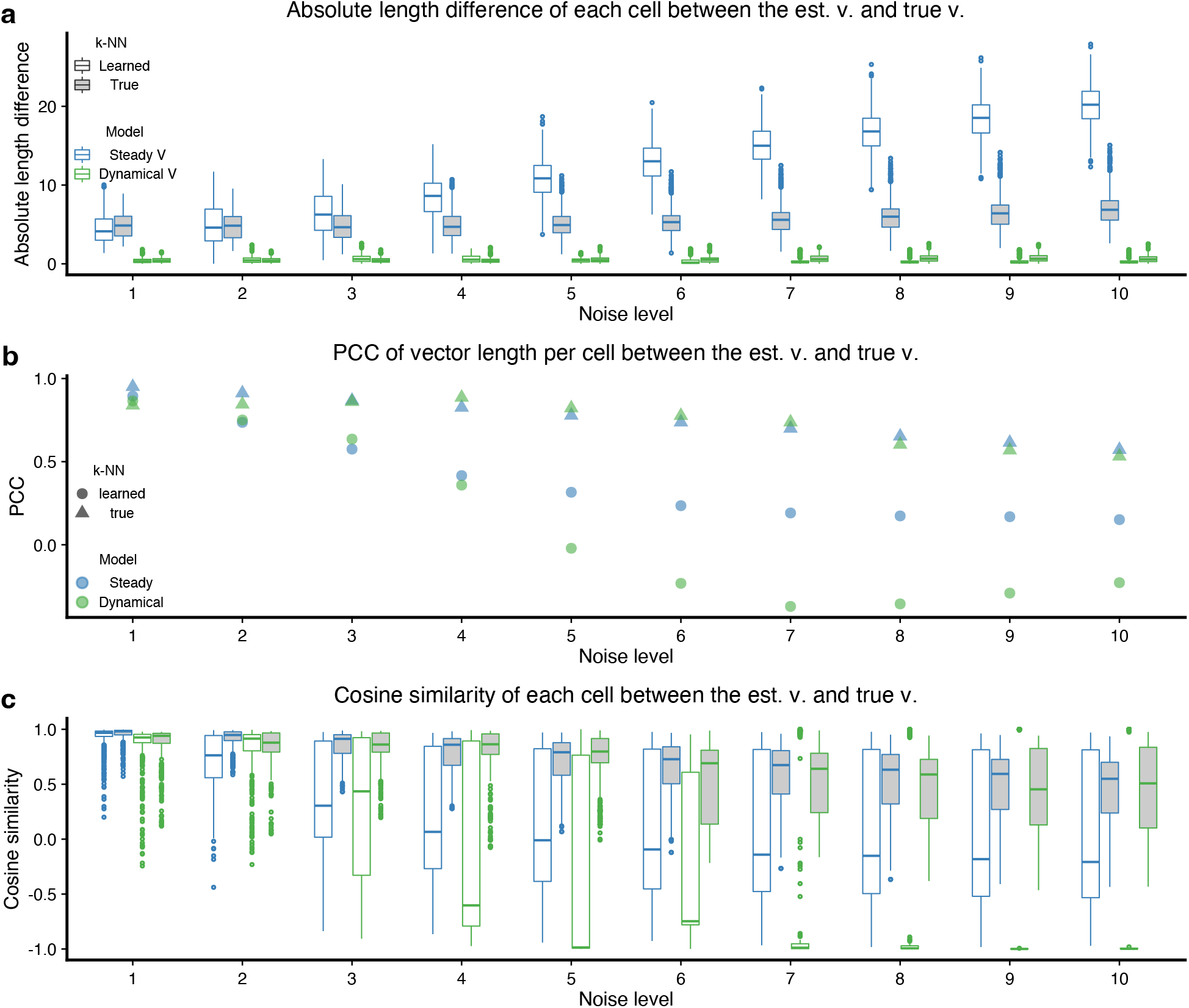
Comprehensive cell-by-cell evaluations of high-dimensional velocity estimations using simulations. **(a)** Boxplots show the absolute vector length difference between the estimated velocities and the true velocities of each cell. **(b)** We show the PCC between the cell-level vector length of estimated velocities and the cell-level vector length of true velocities. **(c)** Boxplots show cosine similarity between the estimated velocities and the true velocities of each cell. All boxes show the left *y*-axis values, and indicate 25th and 75th percentiles. Whiskers extend to the largest values no further than 1.5 × interquartile range (IQR) from these percentiles.

**Supplementary Figure S10.**
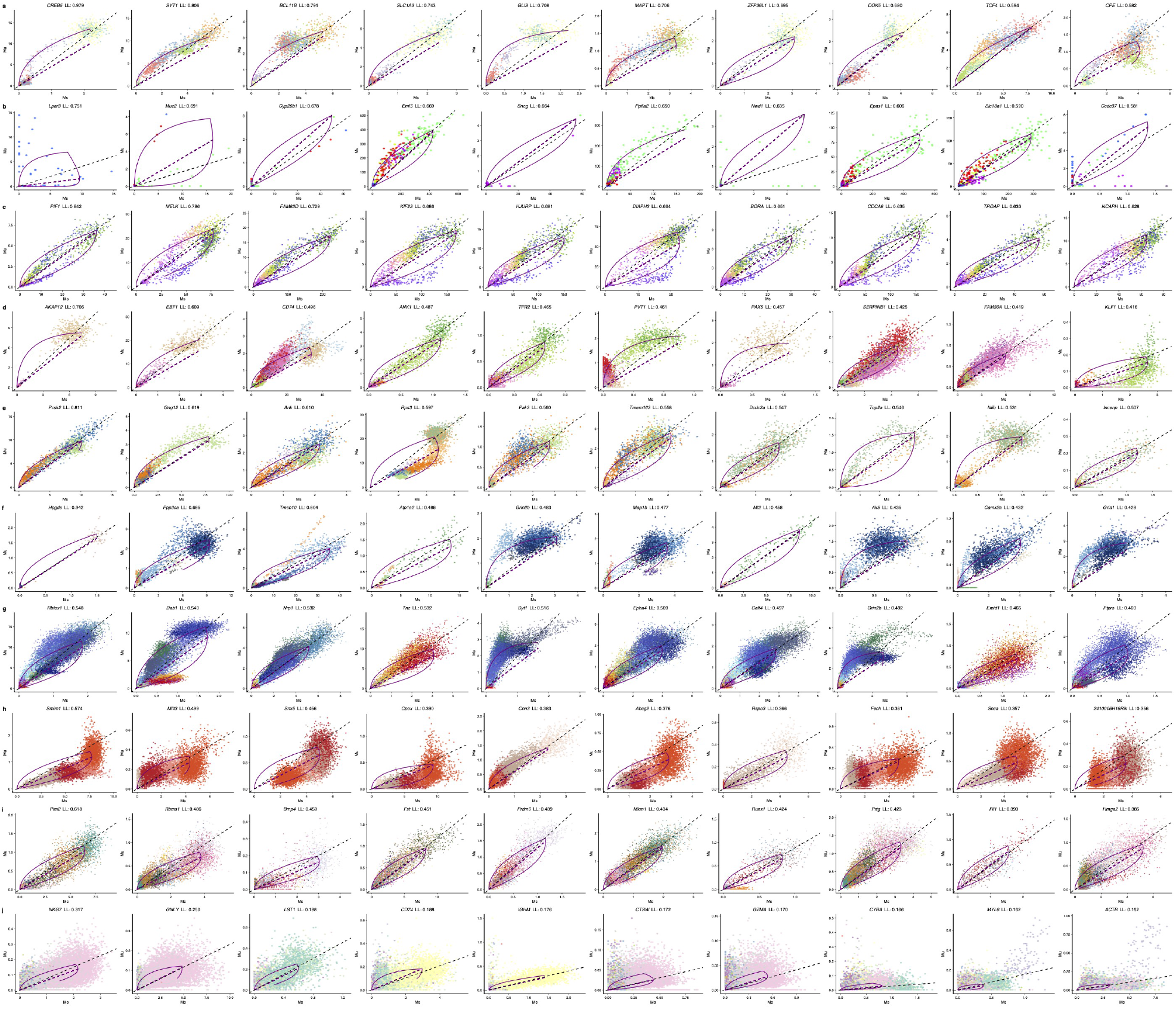
The phase portrait of top 10 genes in the 10 real datasets. Each point represents a cell, colored by cell type or FUCCI pseudotime. Form (a-j), the order of the datasets is listed as in Supplementary Table S1: Forebrain, Chromaffin, FUCCI, Bonemarrow, Dentategyrus Lamanno, Pancreas, Gastrulation erythroid, Dentategyrus Hochgerner, Gastrulation E7.5, and PBMC68k. The top 10 genes ranked by the likelihood from the dynamical model are shown from left to right for each dataset. Purple lines represent the fitted dynamics by the dynamical model, and the black dashed line represents the degradation rate (slope) from the steady-state model. Note that most genes do not show complete up- and down-regulation dynamics.

**Supplementary Figure S11.**
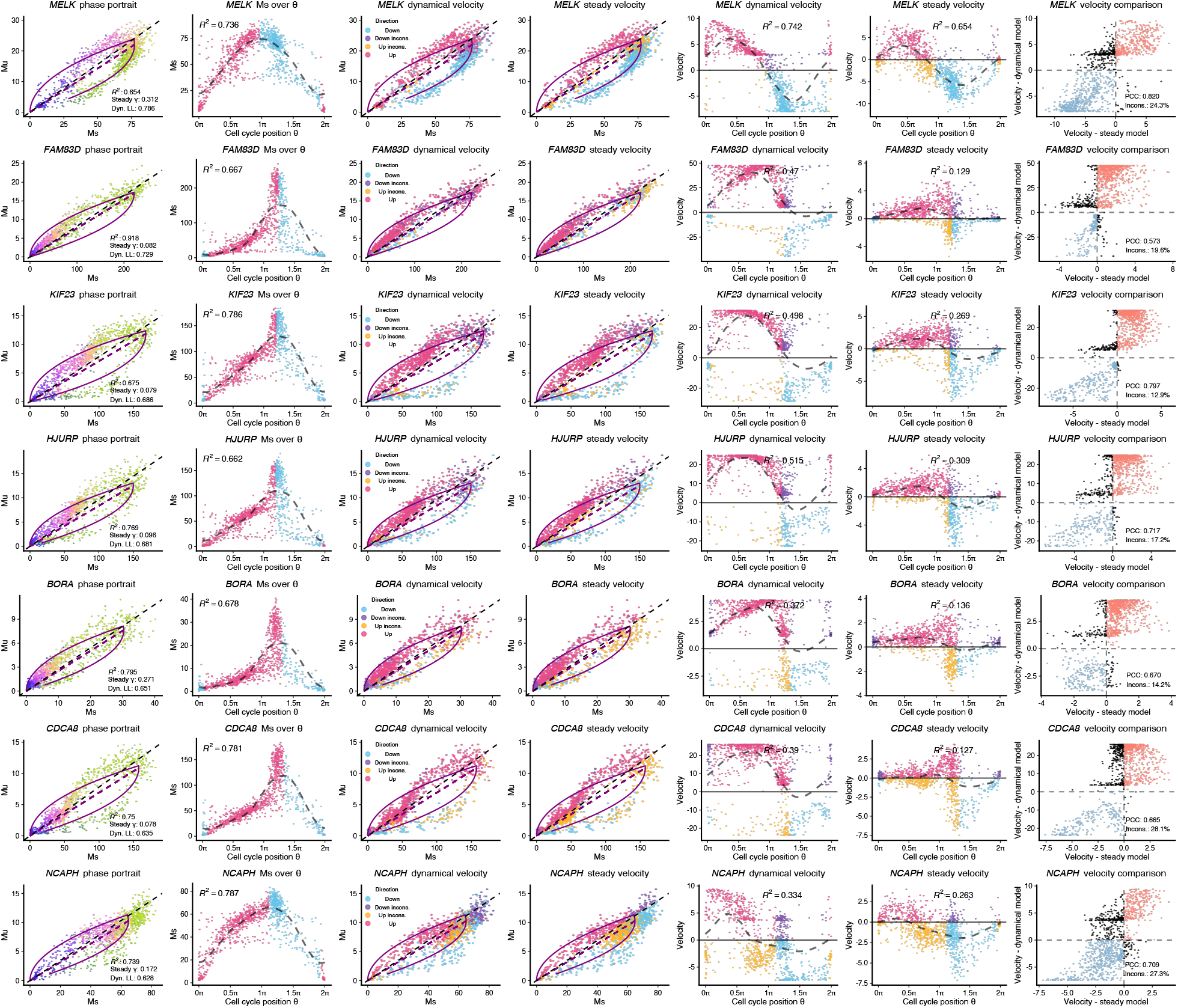
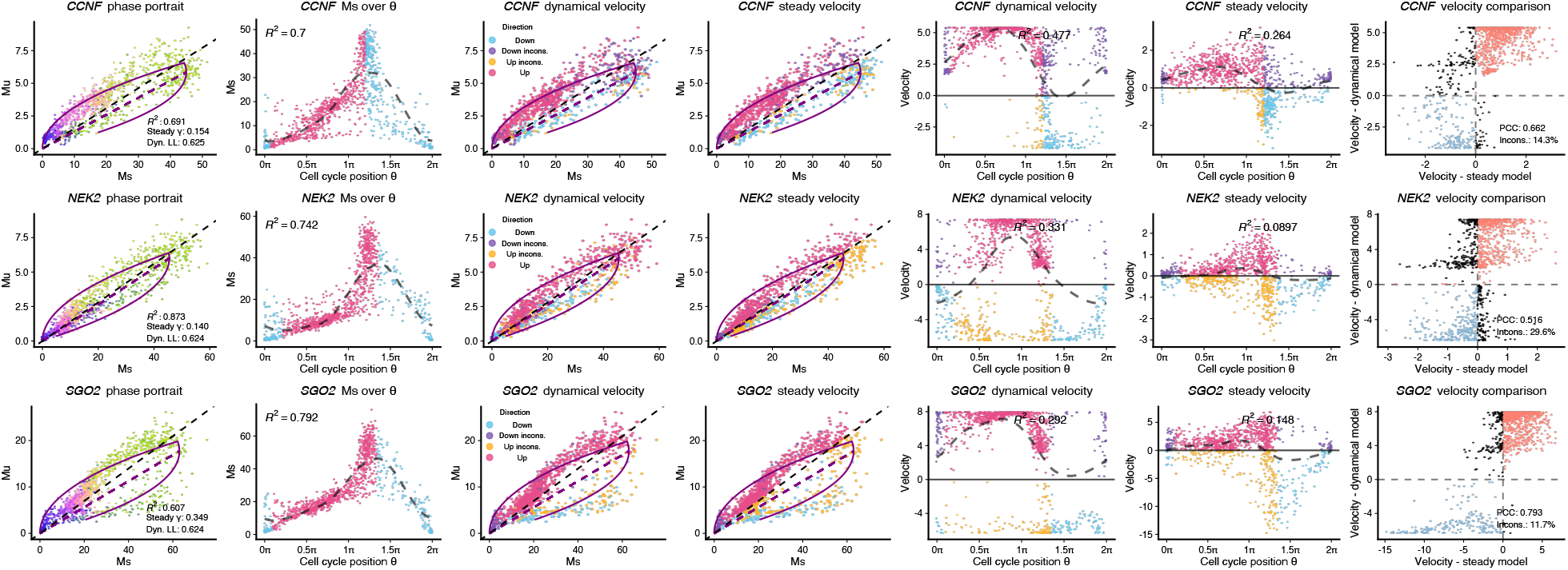
Analyses of the 10 best genes (highest likelihoods) in the FUCCI data. Each row contains a gene, ranked decreasingly by the likelihood. Each point represents a cell in the FUCCI data. From left to right of each row: phase portrait of the gene with points colored by cell cycle position; smoothed expression (Ms) over cell cycle position; phase portrait with points colored by direction comparisons between direction inferred by Ms and dynamical model based velocity estimates; phase portrait with points colored by direction comparisons between direction inferred by Ms and steady-state model based velocity estimates; dynamical model based velocity estimates over cell cycle position; steady-state model based velocity estimates over cell cycle position; comparison of velocity estimates using the steady-state model and the dynamical model.

**Supplementary Figure S12.**
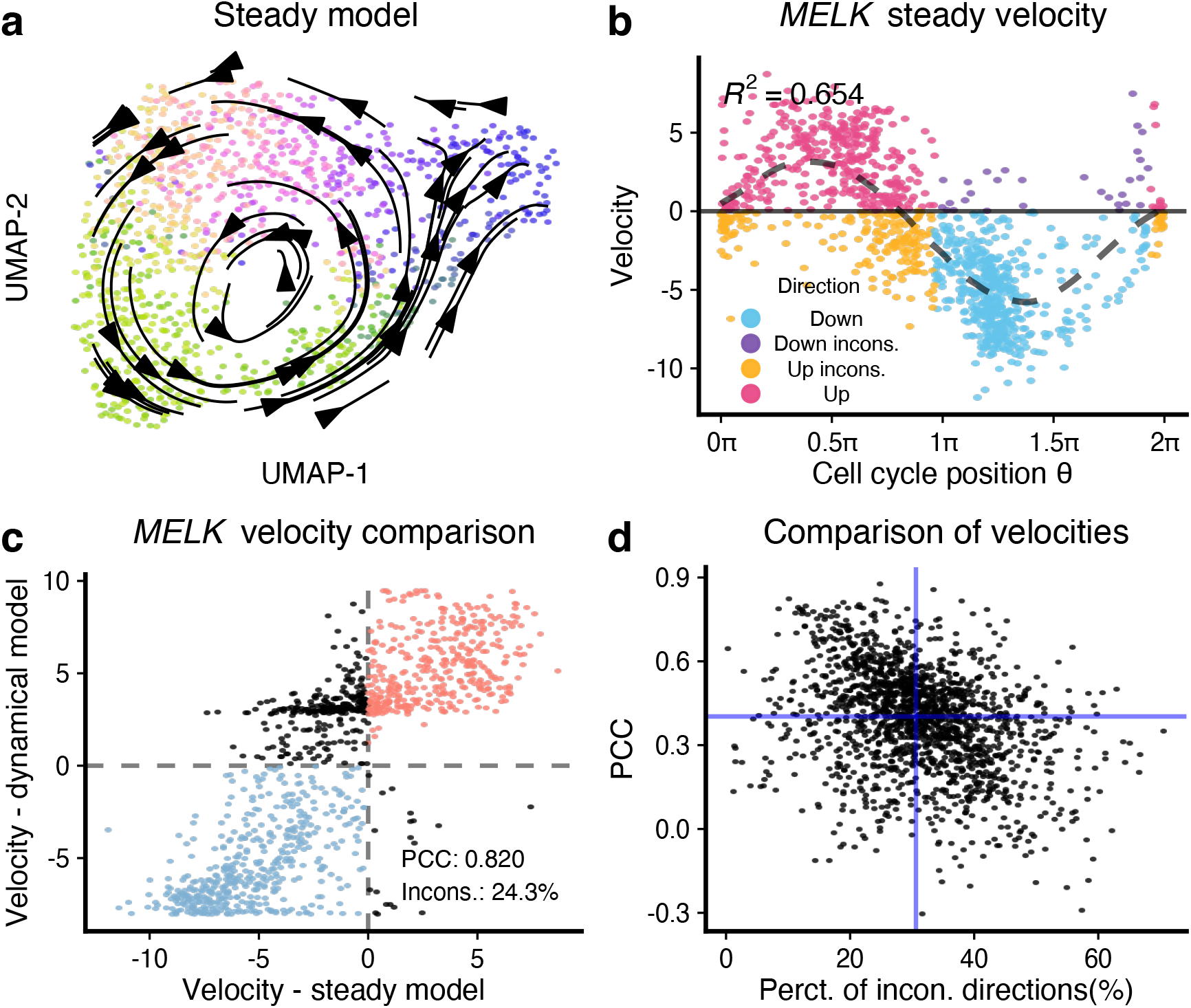
RNA velocity estimation using the steady-state model on the FUCCI data and comparisons of gene-level velocities between steady-state and dynamical model. **(a)** Similar to Figure 6a, but we now use RNA velocity estimates using the steady-state model. RNA velocity vector fields are visualized using the transition probability method on the UMAP embeddings of the FUCCI data. Each point represents a cell and is colored by its FUCCI pseudotime. **(b)** Scatter plot shows the estimated RNA velocity of MELK over FUCCI pseudotime. The signs of velocity estimations are compared to those inferred in Figure 6c, with inconsistent directions colored black. **(c)** Comparison of RNA velocity for gene MELK estimated by steady-state model and dynamical model. About 24.3% of cells, which are colored black, exhibit opposite velocities between the two models. **(d)** Scatter plot shows the PCC and percentage of inconsistent directions between velocities estimated by the steady-state model and dynamical model for all velocity genes in the FUCCI data. The two blue lines give the respective median values, with a PCC median of only about 0.4.

**Supplementary Figure S13.**
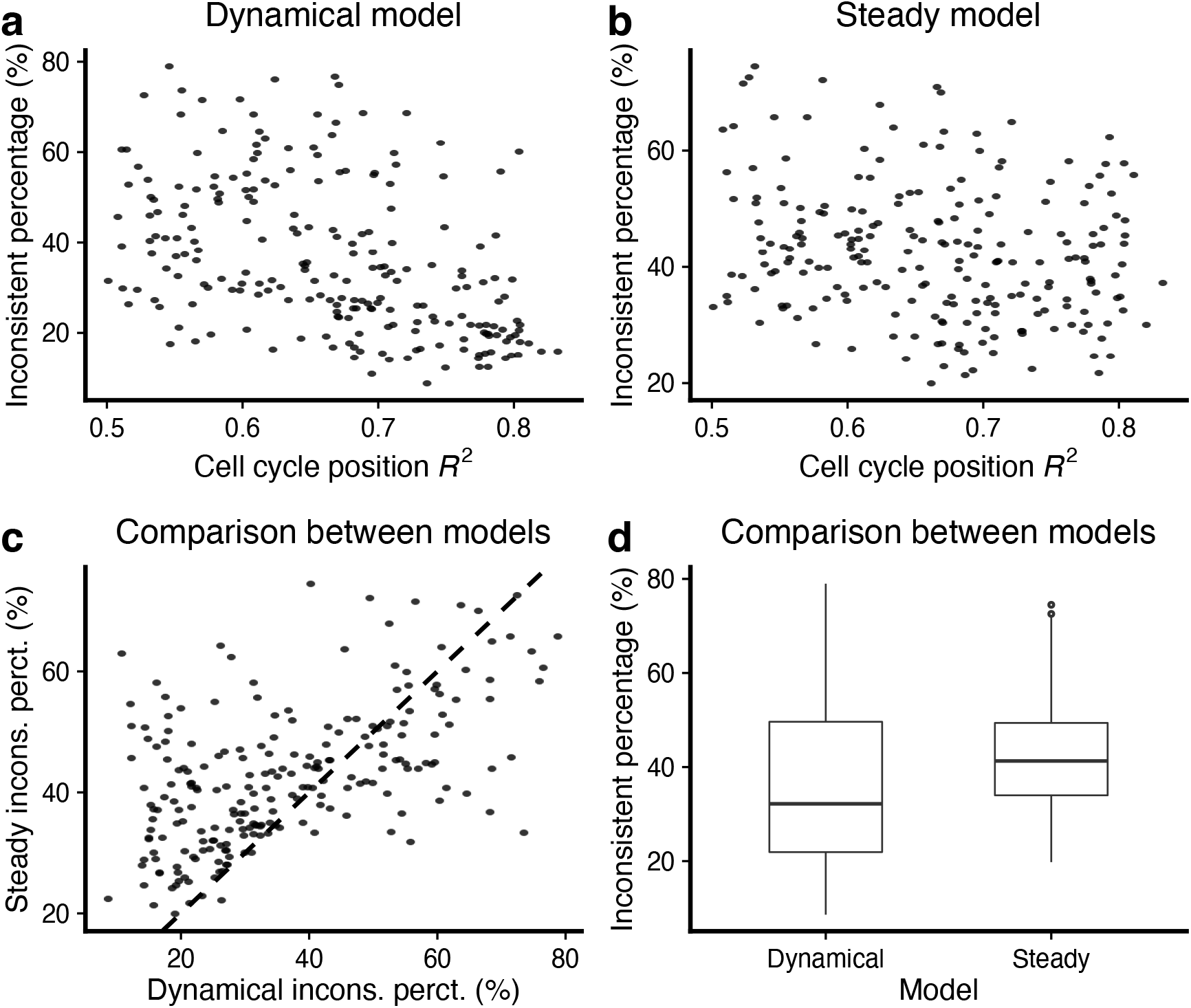
Percentage of cells showing the inconsistent direction of change between fitted model using cell cycle position and gene-specific RNA velocity. Out of 579 cell cycle genes which are also velocity genes, we examine the 224 genes that have a R^2^ of periodic loess over cell cycle position greater than 0.5. **(a)** Scatter plot shows the percentage of cells having the inconsistent direction of change between the fitted model using cell cycle position and dynamical model based gene-specific RNA velocity estimates. **(b)** As (a), but now we use the steady-state model to get RNA velocity estimates. **(c-d)** Comparison of the percentage of cells having the inconsistent direction of change between the steady-state and dynamical model velocity estimates.

**Supplementary Figure S14.**
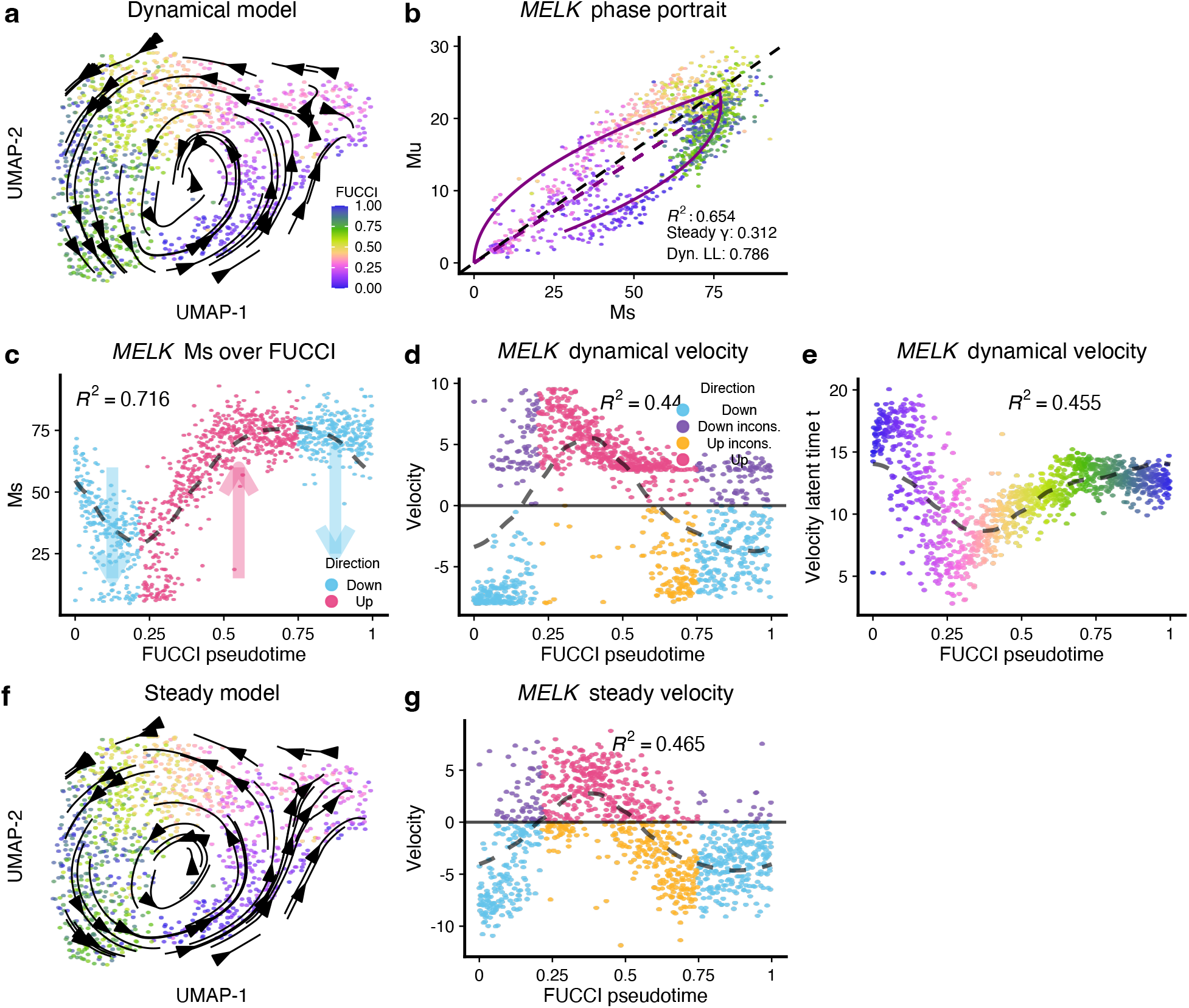
The RNA velocity application on FUCCI dataset (colored by FUCCI pseudotime). **(a-e)** Similar to Figure 6, but now we use FUCCI pseudotime as the “true” trajectory. **(a)** RNA velocity vector field are visualized using the transition probability method on the UMAP embeddings of the FUCCI data. Each point represents a cell and is colored by FUCCI pseudotime. **(b)** Phase portrait of gene *MELK*, of which the likelihood is the highest among all velocity genes inferred by the dynamical model. The purple lines represent the dynamics inferred by the dynamical model. **(c)** Scatter plot shows smoothed expression of *MELK* over FUCCI pseudotime. The dashed line is the fitted line by periodic loess (Methods). The expected direction of change is inferred on the fitted loess line and visualized by colors. **(d)** Scatter plot shows the estimated RNA velocity of *MELK* over FUCCI pseudotime. The signs of velocity estimates are compared to those inferred in (c), with inconsistent directions colored as black. **(e)** The level of agreement between the velocity latent time for *MELK* and the FUCCI pseudotime is lower than between the velocity latent time and the cell cycle position. **(f)** As (a), but we use the steady-state model velocity estimates. **(g)** As (d), but we use the steady-state model velocity estimates.

**Supplementary Figure S15.**
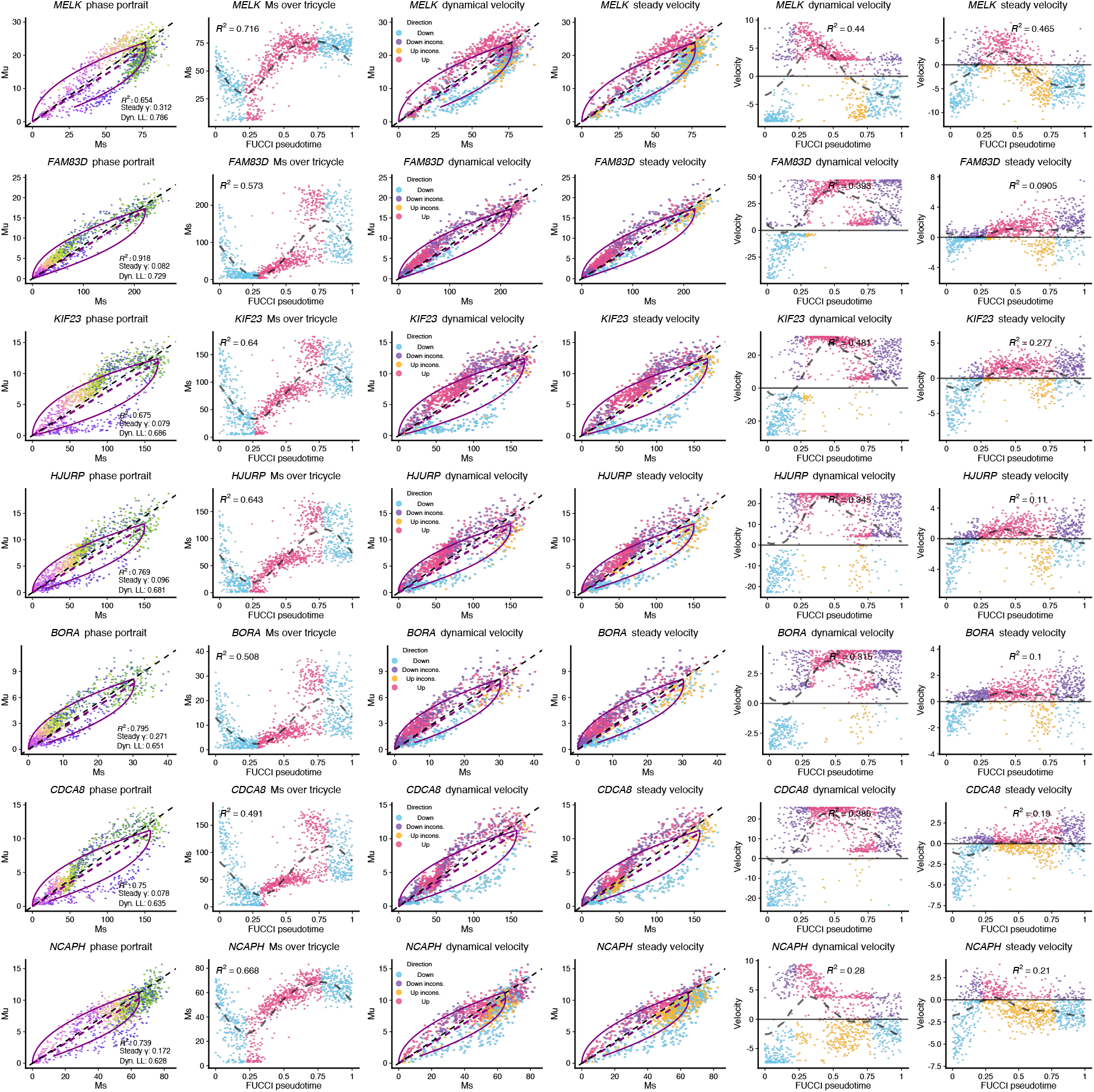
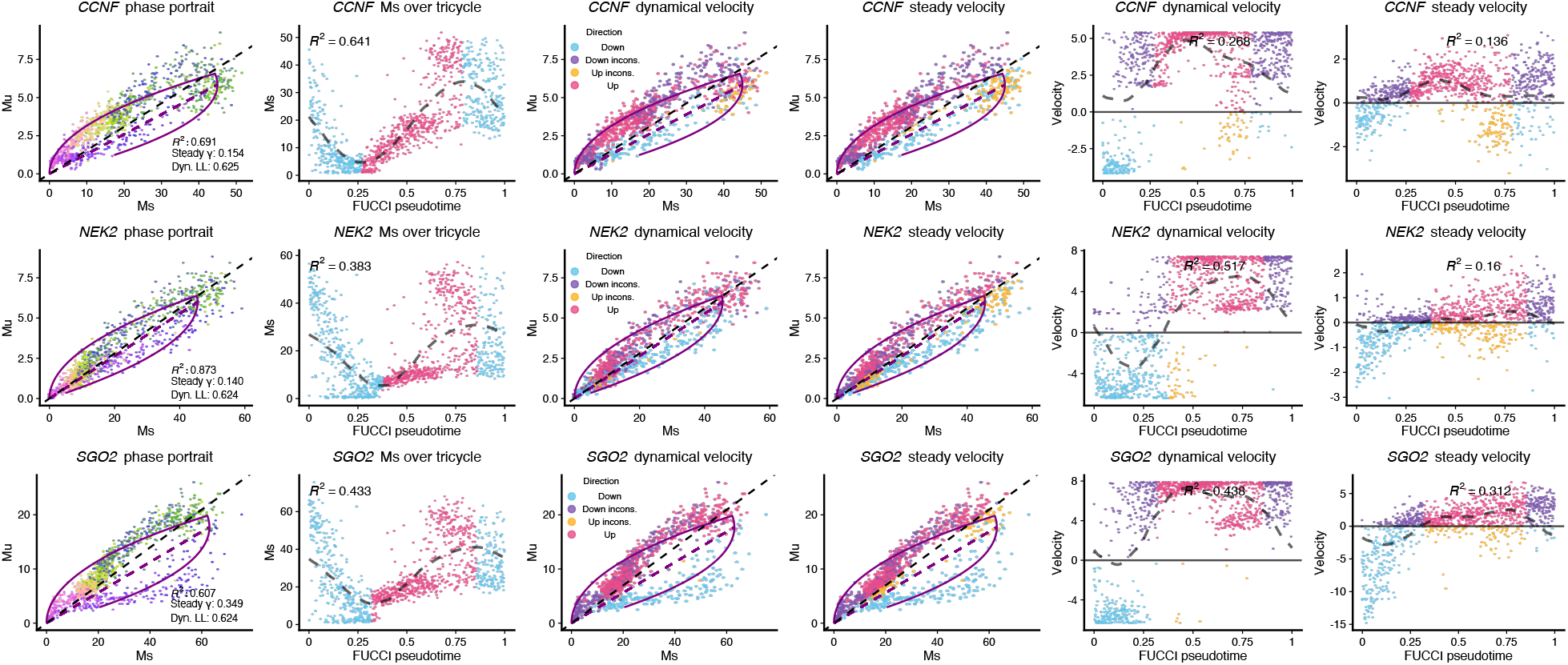
Analyses of the 10 best genes (highest likelihoods) in the FUCCI data (using FUCCI pseudotime). This figure is similar to Supplementary Figure S11, but now we use FUCCI pseudotime as the “true” trajectory. Each row contains a gene, ranked decreasingly by the likelihood. Each point represents a cell in the FUCCI data. From left to right of each row: phase portrait of the gene with points colored by FUCCI pseudotime; smoothed expression (Ms) over FUCCI pseudotime; phase portrait with points colored by direction comparisons between direction inferred by Ms and dynamical model based velocity estimates; phase portrait with points colored by direction comparisons between direction inferred by Ms and steady-state model based velocity estimates; the dynamical model based velocity estimates over FUCCI pseudotime; the steady-state model based velocity estimates over FUCCI pseudotime.

### 2 SUPPLEMENTARY TABLES

**Supplementary Table S1.** Datasets.

| Dataset | Species | Platform | Data Access | Reference |
| --- | --- | --- | --- | --- |
| Forebrain | Human | 10x | scvelo | La Manno et al. (2018) |
| Chromaffin | Mouse | SMART-seq2 | velocyto | Furlan et al. (2017) and La Manno et al. (2018) |
| FUCCI | Human | SMART-seq2 | GSE146773 | Mahdessian et al. (2021) |
| Bonemarrow | Human | 10x | scvelo | Setty et al. (2019) |
| Dentategyrus LaManno | Mouse | 10x | scvelo | Hochgerner et al. (2018) and La Manno et al. (2018) |
| Pancreas | Mouse | 10x | scvelo | Bastidas-Ponce et al. (2019) and Bergen et al. (2020) |
| Gastrulation erythroid | Mouse | 10x | scvelo | Pijuan-Sala et al. (2019) |
| Dentategyrus Hochgerner | Mouse | 10x | scvelo | Hochgerner et al. (2018) and Bergen et al. (2020) |
| Gastrulation E7.5 | Mouse | 10x | scvelo | Pijuan-Sala et al. (2019) |
| PBMC68k | Human | 10x | scvelo | GXY Zheng et al. (2017) |

### 3 SUPPLEMENTARY NOTES

#### 3.1 The differences between implementations of RNA velocity analysis

Apart from the fact that the scVelo implements several models for high dimensional RNA velocity estimations and velocyto (Python and R packages) only implements a steady-state model, the default workflows of the three packages differ in many steps. Here, we discuss some key differences that we think are important.

In the high-dimensional gene-level RNA velocity estimation step, all implementations smooth (also referred to as imputation) the spliced and unspliced counts using a k-NN graph, but the details differ. The scVelo package builds a k-NN graph using the Euclidean distance between cells in a low-dimensional PCA space (derived from the spliced matrix), followed by a weighted smoothing step with weights computed as in McInnes et al. (2018). Both parameter estimation and velocity computation use these smoothed values. The python implementation of velocyto builds a k-NN graph using Euclidean distance in a low dimensional PCA space (using a different k-NN implementation from scVelo) but then performs unweighted smoothing. Both parameter estimation and velocity computation use these smoothed values. In contrast, the R implementation of velocyto uses an unweighted k-NN graph using the correlation distance on the high-dimensional spliced counts matrix. Neighboring cells are aggregated into pseudo cells, and the quantile regression line is fitted on these pseudo cells to get the estimated degradation rate. After calculating the degradation rate, velocities are computed for each gene by plugging in the unspliced and the spliced counts (prior to forming pseudo-cells) in Equation 2; a substantial deviation from the Python implementation.

Another big difference lies in choosing a number *K* of the k-NN graph. The default parameters used in the scVelo package are *k* = 30 neighbors. For velocyto, there are no defaults for the *k*, but the analyses presented in La Manno et al. (2018) sometimes use huge values (e.g., *k* = 550 for a forebrain dataset with less than 2,000 cells). As the *k* is the most important parameter for all k-NN graphs, which is central to the high-dimensional RNA velocity estimation step, the choice of *k* might substantially affect the visualized low-dimensional vector field in the end. As an example, we compare the RNA velocity vector field of the Forebrain data by using *k* = 30 and *k* = 550 for both steady-state and dynamical models (we use scVelo implementation for both models here to avoid differences caused by other steps). Comparing Supplementary Figure S16a to Supplementary Figure S16b, we see that the use of a larger *k* in the steady-state model seems to give us a more smooth vector field. Nevertheless, we observe a fairly striking difference for the dynamical model where the low-dimensional vectors of neuroblast cells (purple and red cells) showing opposite direction when using *k* = 30 (Supplementary Figure S16c to using *k* = 550 (Supplementary Figure S16d). Given there is no guide on choosing the *k* in a new dataset, we have consistently used the *k* = 30, the default in the scVelo package.

There are differences between implementation and implementation in the low-dimensional vector field visualization step. A prominent example is that another k-NN graph is constructed using the lowdimensional embedding and used in velocyto to get the Pearson correlation matrix (the cosine similarity matrix in scVelo). In contrast, scVelo uses the same k-NN graph for processing (smoothing) and lowdimensional velocity vector field visualization. For velocyto, the resulting transition probability matrix would be embedding-dependent, and the embedding-dependent property of a transition probability matrix seems counterintuitive.

The three implementations diverge sharply in critical steps without examining how these choices affect the resulting output, including both high-dimensional velocities and low-dimensional vector fields. We won’t pursue the issue further, but we only use the scVelo package for both the steady-state and dynamical models to avoid implementation differences.

**Supplementary Figure S16.**
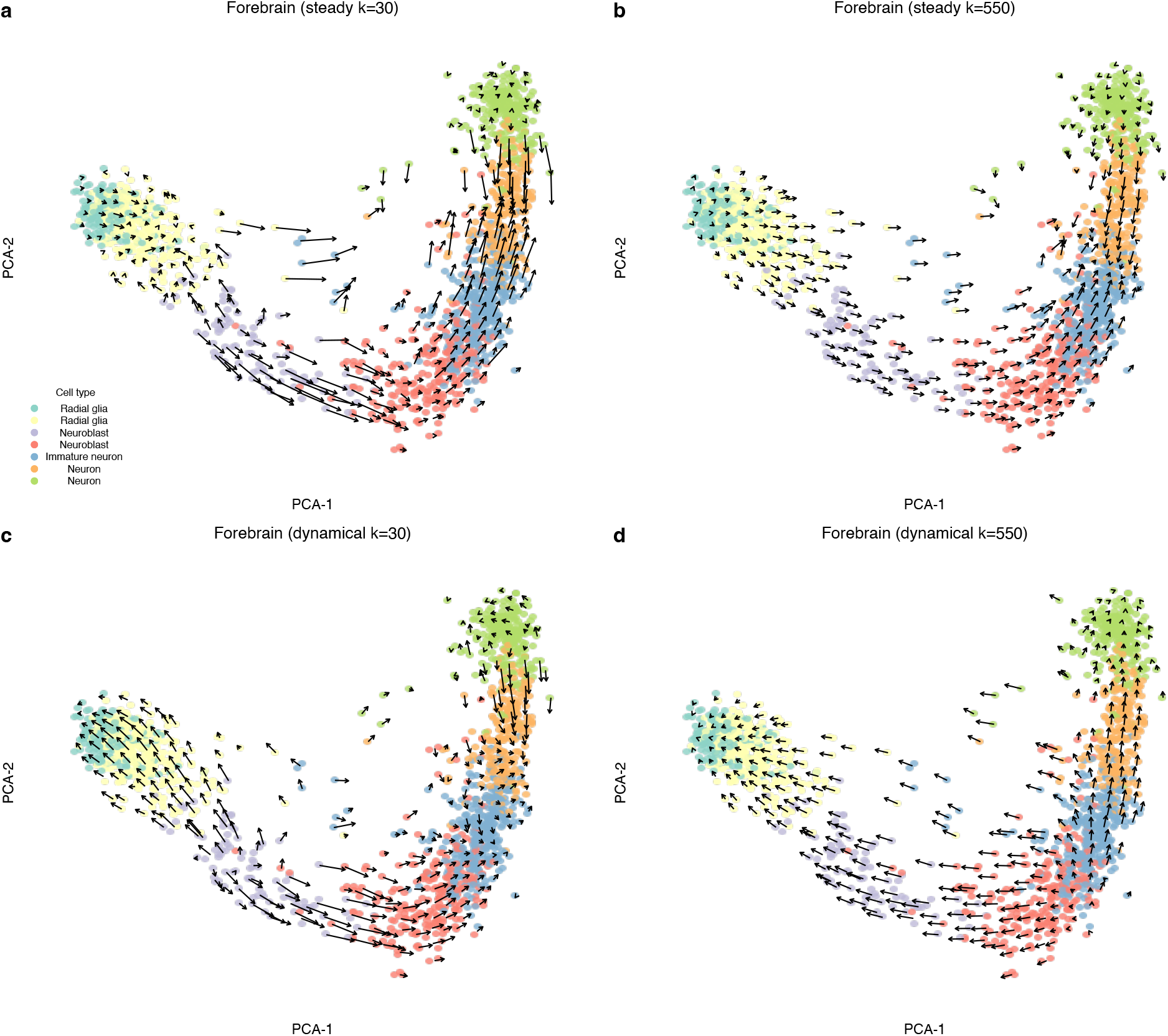
Embedded RNA velocity of the Forebrain data using different number of *K* for the k-NN graph. **(a)** Embedded steady-state model based RNA velocity using the transition probability method of the Forebrain dataset when *k* = 30 for the k-NN graph. **(b)** As (b), but now we use *k* = 550, which was used by La Manno et al. (2018). **(c-d)** As (a-b), but now we use dynamical model to estimate RNA velocity instead.

#### 3.2 Visualization of 2d vector fields

To avoid overplotting, we usually summarize vectors in a 2D space, which can hide local detail. One approach is to grid the 2d embedding and compute the average vectors at each grid location. An alternative is the streamline plot which has multiple distinct implementations (The Matplotlib development team, 2022; Campitelli,2022). Supplementary Figure S17a shows a streamline plot of a forebrain dataset previously discussed in the literature on RNA velocity (La Manno et al., 2018; Gorin et al., 2022). The gridding display of the same vector field produces a very different impression (Supplementary Figure S17b). If we examine the embedding by the left, middle, and right parts, the streamline plot hides that the length of vectors on the left is much shorter than the length of vectors in the middle.

**Supplementary Figure S17.**
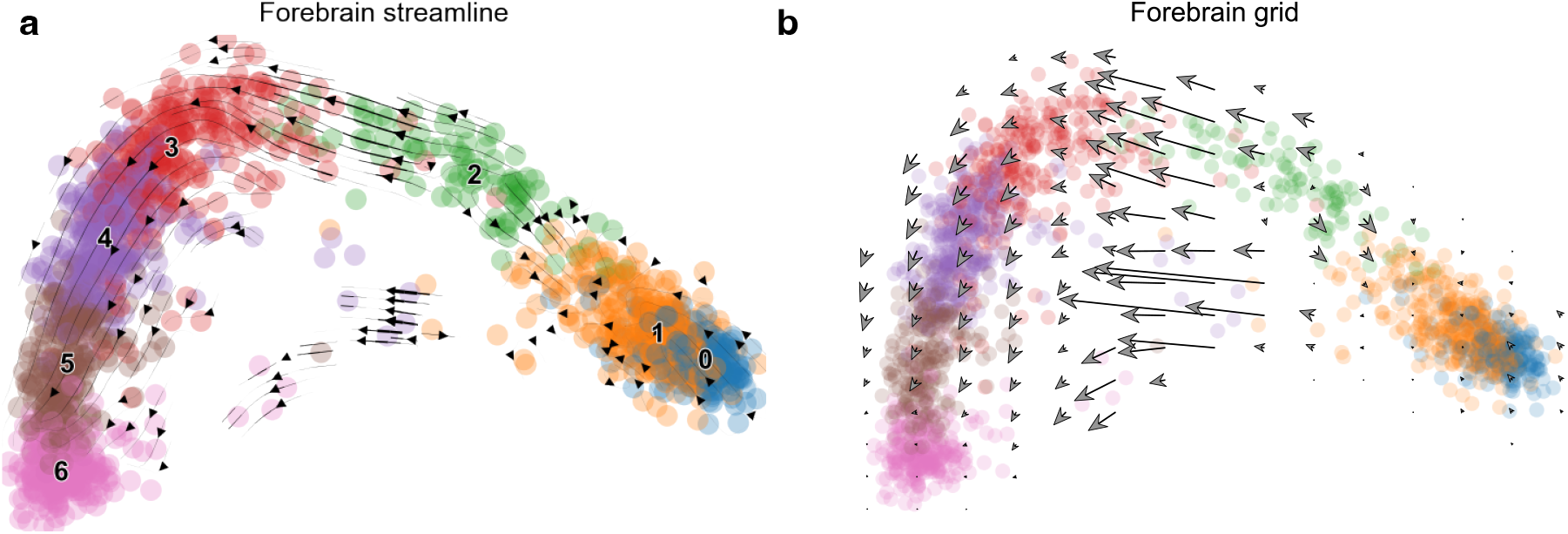
The impact of different visualization approaches. We run scVelo on the Forebrain data and visualize the exact same vector field using two visualization functions pl.velocity_embedding_stream and pl.velocity_embedding_grid in scVelo, which give us quite different impression how the vector field looks like. **(a)** The vector field is visualized by streamline plot (The Matplotlib development team,2022). We use all default parameters for the pl.velocity_embedding_stream function. **(b)** The vector field is visualized by gridding and kernel smoothing. We had to decrease the resolution (density=0.3), increase the arrow size (arrow_size=4), and scale up the arrow length (arrow_length=3) to make arrows visible. Note that in the middle part, we see some long arrows, while the arrows on the other parts are fairly short.

In addition to the choice of visualization approaches, options such as resolution also matter. Not only does the different selection of resolution change the aesthetic impression, but it also affects the interpretation of the vector fields (Supplementary Figure S18), especially locally for some regions in the embedding, such as the top right part of the pancreas data.

These illustrate how the qualitative impression of a vector field depends on the visualization method. As the qualitative impression is usually subjective and highly parameter-dependent, we will not pursue this critical point further. But we strongly recommend using the same visualization tool with choices of appropriate parameters to compare different vector fields. We tend to favor the gridding approach, as it more faithfully depicts local vector fields than the streamline plot and is easier to reason.

**Supplementary Figure S18.**
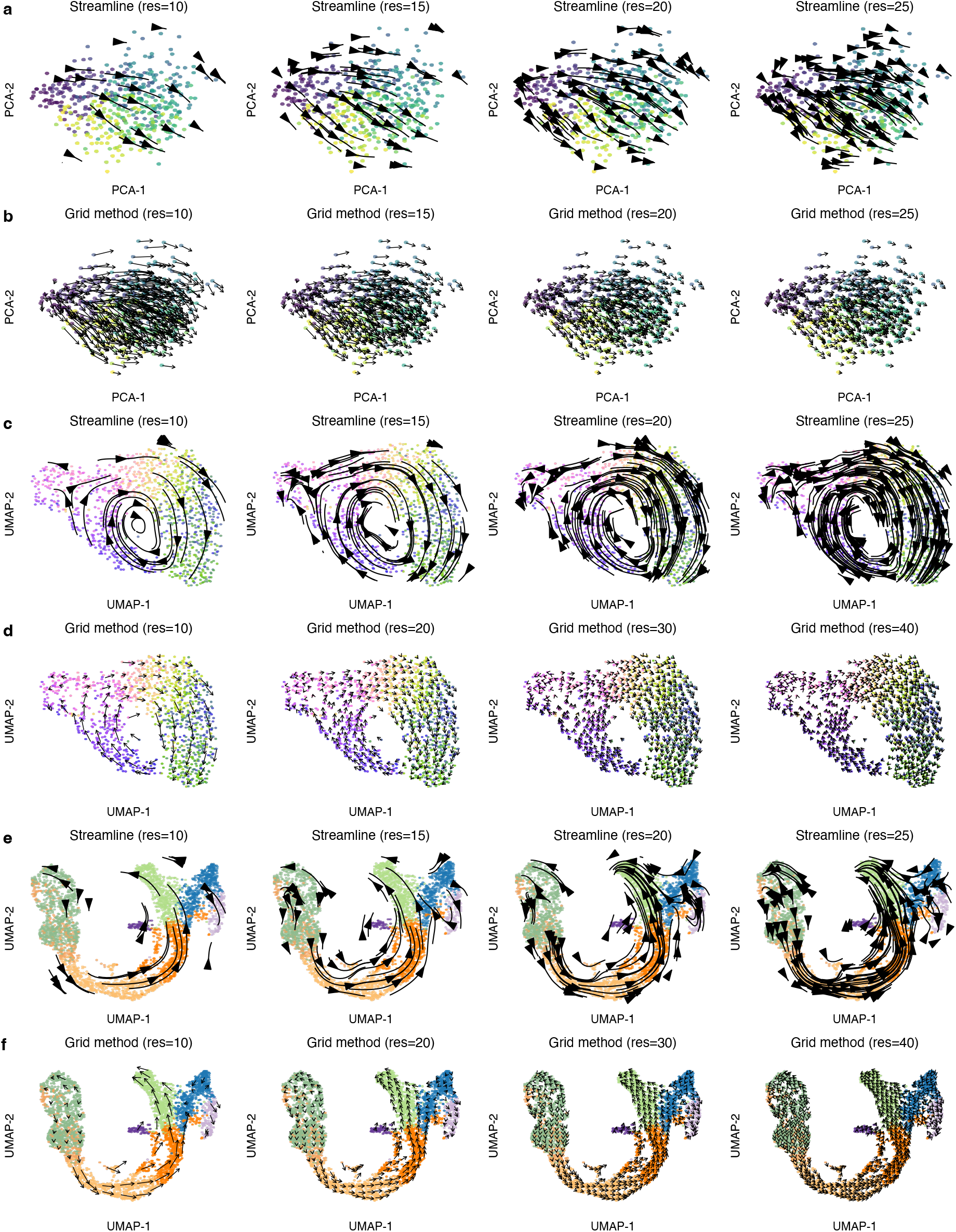
The choice of resolution could affect how the RNA velocity vector field looks. We use three datasets to show that the choice of resolution could affect how the RNA velocity vector field looks like: simulated data in (a-b); FUCCI data in (c-d); pancreas data in (d-e). For those datasets, we used the dynamical model RNA velocity estimates. We use the streamline method to visualize the velocity vector field in (a), (c), and (e), while we use the grid method in (b), (d), and (f). From left to right, we increase the level of resolution. Note that how good the vector field looks depends on the subjectively optimal choice of resolution level, which is difficult to decide and varies across datasets.

## Footnotes

1 There are many different terminologies for the RNA velocity vector field visualization. For example, Bergen et al. (2020) used “project” to describe transforming the gene-level RNA velocity into an embedding. We use “map” to describe the process of transforming the high-dimensional gene-level RNA velocities into the low-dimensional cell-specific vectors that can be visualized in the embedding space. And we reserve the word “project” to describe a mathematical projection onto a linear subspace. A large number of cells in most datasets implies that the final visualizations depict summa-rized/smoothed vector fields from the low-dimensional cell-specific vectors to avoid overplotting. Hence, we use “visualized” to describe the final visualized vector field.

